# Odor-based real-time detection of pests on maize plants

**DOI:** 10.1101/2024.07.29.605549

**Authors:** Marine Mamin, Carla C. M. Arce, Gregory Röder, Arooran Kanagendran, Thomas Degen, Emmanuel Defossez, Sergio Rasmann, Terunobu Akiyama, Kosuke Minami, Genki Yoshikawa, Felipe Lopez-Hilfiker, Priyanka Bansal, Luca Cappellin, Yunhe Li, Ted C. J. Turlings

**Author notes:** These authors contributed equally to this work (Marine Mamin, Carla C. M. Arce).

## Abstract

Early detection of crop pests and diseases can enable timely, targeted interventions, and help reduce pesticide use. Plants under biotic stress are known to rapidly emit characteristic blends of volatile compounds that could potentially serve as early and attacker-specific cues for precise pest monitoring. Here, we evaluated the feasibility of this approach using two complementary, state-of-the-art sensing technologies: a handheld nanomechanical membrane-based sensor array and chemical ionization time-of-flight mass spectrometry. Under laboratory conditions, with enclosed headspace sampling, both technologies readily distinguished undamaged maize plants from plants infested by caterpillars or infected with a fungal pathogen. Under semi-controlled outdoor open-air conditions, where volatile concentrations were strongly diluted, the membrane-based sensor no longer retained discriminatory power, whereas mass spectrometry predicted herbivory status with more than 90% accuracy using one-second measurements. Finally, in an initial field trial based on simulated herbivory, a compact, field-deployable, real-time mass spectrometer distinguished damaged from undamaged maize plants with highly encouraging performance under real field conditions. Together, these results demonstrate the potential of odor-based detection of pest attacks in maize and identify real-time mass spectrometry as a promising tool for crop monitoring, while pinpointing challenges that remain to be addressed for translation to practical field applications.

## Introduction

Current agricultural practices depend heavily on synthetic agrochemicals, a reliance widely acknowledged as unsustainable and a major contributor to environmental pollution, habitat degradation, biodiversity loss, and climate change (Poore and Nemecek, 2018; Tang et al., 2021). Reducing agrochemical use while safeguarding yields is therefore an essential focus of current agricultural research (Möhring et al., 2023). Despite decades of innovative control efforts, pests and diseases still cause up to 40% of potential crop losses (Savary et al., 2019), further underscoring the need for more effective and sustainable approaches to plant protection (Godfray et al., 2010; Pretty et al., 2018; Tang et al., 2025).

Early detection of pests and diseases through real-time monitoring can greatly facilitate timely and precise interventions, reducing pesticide use while maintaining crop protection (Getahun et al., 2024; Godavari et al., 2024; Ye et al., 2025). Precision agriculture has advanced several non-destructive monitoring techniques, including methods that directly detect attackers (e.g. acoustic sensors for insect activity) and those that monitor changes in plant phenotypes through imaging technologies (Bukhamsin et al., 2025; He et al., 2023; Singh et al., 2020). Despite significant progress, key challenges remain, particularly detecting attacks at early stages and discriminating between multiple stressors under field conditions (Chauhan et al., 2025).

Inducible plant volatiles represent promising early warning signals of pest and disease presence (Cui et al., 2018; Tholl et al., 2021; Coatsworth et al., 2022; MacDougall et al., 2022; Fundurulic et al., 2023; Gan et al., 2023; Zheng et al., 2023). In response to arthropod herbivory or infection by pathogenic agents, plants swiftly emit characteristic blends of volatiles, often just within hours of attack (Turlings et al., 1998b; Heil, 2014; Schuman et al., 2016; Sharifi et al., 2018). These emissions are shaped by plant recognition of elicitors produced by the attackers, such as pathogen-derived molecules or compounds present in oral and egg secretions of herbivores (Alborn et al., 1997; Boller and Felix, 2009; Hilker and Fatouros, 2015; Jones et al., 2022). Because different elicitors activate distinct signaling cascades, the resulting odor blends can be highly specific (Erb and Reymond, 2019; Jones et al., 2022). This specificity is exemplified by the ability of natural enemies to locate plants carrying their preferred insect prey (De Moraes et al., 1998; Allmann and Baldwin, 2010). Thus, inducible plant volatiles can serve as attacker-specific “chemical fingerprints” that can potentially be exploited for both early and precise crop surveillance (Turlings et al., 1998b).

Two main strategies have been proposed to exploit plant volatiles for monitoring crop health. Static systems, such as leaf-wearable sensors, could continuously track individual plants (Li et al., 2021; Lee et al., 2024), while mobile systems could sample volatiles across fields, potentially through autonomous platforms (Geckeler et al., 2023). A mobile solution, however, requires odor sensing technologies capable of fast, real-time measurements, robust to environmental fluctuations and with sufficient sensitivity to detect volatiles in open air, where they are diluted, variable, and intermingled with background odors.

Different portable electronic gas sensors have been developed for rapid onsite odor detection, using arrays of sensing elements whose chemical or physical properties change upon interaction with volatiles. These changes are transduced into electronic or optical signals and analyzed with pattern recognition algorithms to discriminate odor profiles (Li et al., 2019; Ivaskovic et al., 2021; MacDougall et al., 2022; Zheng and Zhang, 2022; Wesoły et al., 2023; Zheng et al., 2023; Chen et al., 2024). While several of these small systems have demonstrated the ability to detect volatile signatures associated with pest or pathogen attack in agriculturally relevant plant species, most studies have been performed under laboratory or greenhouse conditions, and using enclosed sampling setups, minimizing dilution and background interference (Cui et al., 2018; Tholl et al., 2021; Zheng and Zhang, 2022; Fundurulic et al., 2023). In addition, while much larger, high-performance instruments can provide valuable capabilities for real-time odor detection. In particular, Chemical Ionization Time-Of-Flight Mass Spectrometry (CI-TOF-MS) is highly performant for online measurement of trace volatiles in open-air and complex environments, as showcased by its use in atmospheric field studies (Hutterli et al., 2022).

We investigated here the feasibility of real-time pest and disease detection based on volatiles emissions of maize (*Zea mays*), one of the most widely cultivated crops worldwide. Experiments progressed from laboratory assays using a dynamic headspace sampling system to open-air measurements in semi-controlled outdoor conditions and finally initial field measurements. We applied two complementary and existing sensing technologies. The first was a handheld module incorporating Membrane-type Surface Stress Sensors (MSS – NIMS), which detect volatile-receptor interactions through membrane deformation that generates piezoresistive signals (Yoshikawa et al., 2011, 2012; Minami et al., 2022). This compact electronic sensor has demonstrated potential for food, environmental, and medical diagnostics (Imamura et al., 2016; Lang et al., 2016; Minami et al., 2023; Saeki et al., 2024). The second technology was CI-TOF-MS. Laboratory and open-air assays were conducted with an instrument based on proton transfer reaction (PTR-TOF; Vocus S – Tofwerk AG) (Krechmer et al., 2018), an ionization method previously used to study dynamic stress responses in plants (Niederbacher et al., 2015; Majchrzak et al., 2020; Tholl et al., 2021; Waterman et al., 2025). For field measurements we used an ultra-portable TOF instrument (Vocus C – Tofwerk AG) Dobrecevich et al., 2026), configured with benzene cation ionization (Puttu e tal., 2025; Kim et al., 2016, Riva et al., 2024). Laboratory studies homed in on volatiles induced by two lepidopteran pests of maize, *Spodoptera exigua* and *S. frugiperda,* and the fungal pathogen *Colletotrichum graminicola*, whereas subsequent experiments focused on herbivory by *Spodoptera* caterpillars, given their agricultural significance (Kartakis et al., 2025). This study provides an initial assessment of the feasibility of odor-based pest and disease detection in maize using available technologies, offering a foundation for future efforts to integrate this approach into agriculture to optimize and reduce agrochemical use.

## Results

### 1) Laboratory measurements

#### 1.1) Volatile profiling of maize plants using GC–MS

Volatile emissions from undamaged maize plants and plants infected with *C. graminicola* or infested with *S. frugiperda* or *S. exigua* were first characterized under laboratory conditions using dynamic headspace sampling and GC-MS analysis (Fig. S1). Two commercial hybrids were examined (var. *Delprim* and *Aventicum*). Nonmetric multidimensional scaling (NMDS) revealed clear separation among volatile profiles of undamaged, fungus-infected, and caterpillar-infested plants for both varieties (Fig. 1A; PERMANOVA: Delprim: F_3,26_ = 28.865, p = 0.001; Aventicum: F_3,28_ = 17.662, p = 0.001).

**Fig. 1.**
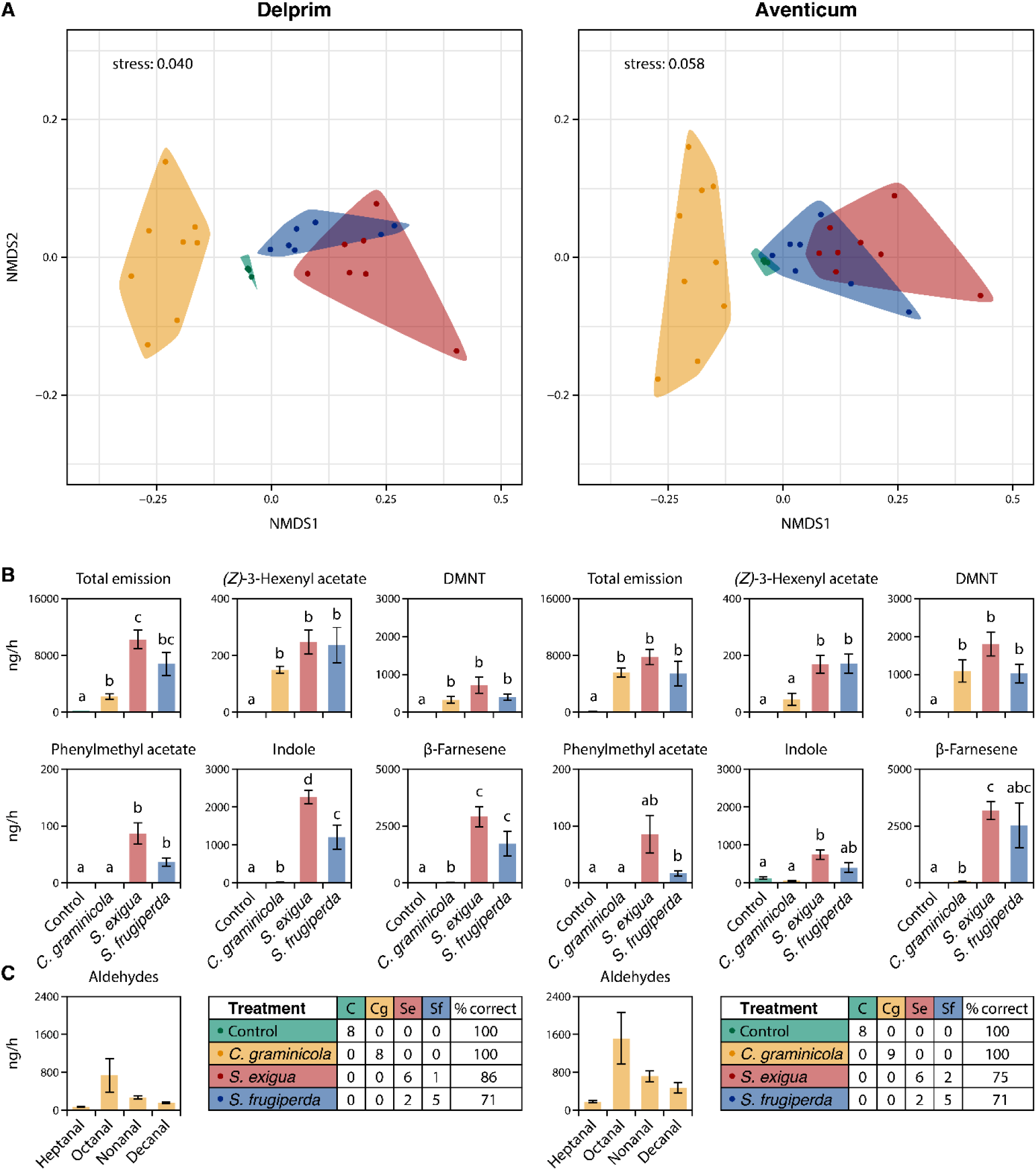
Characterization of the volatile profiles of maize plants infested with different pests. Volatile emissions were collected under laboratory conditions using classical push–pull headspace sampling and analyzed by GC-MS. Undamaged maize plants (green) of the varieties *Delprim* and *Aventicum* were compared with plants infested by a fungal pathogen (*Colletotrichum graminicola*; yellow) or by caterpillars (*Spodoptera exigua*, red; *Spodoptera frugiperda*, blue). **(A)** NMDS ordination illustrating differences in overall volatile profiles among treatments. **(B)** Emission rates (ng h⁻¹; mean ± s.e.) of selected major volatile compounds. Different letters indicate significant pairwise differences after FDR correction; full results in Supplementary Information. **(C)** Confusion matrix summarizing the performance of a Random Forest classification model in assigning volatile profiles to the correct treatment; prediction accuracy was estimated using the model’s internal out-of-bag samples.

Undamaged plants constitutively released only small amounts of linalool in *Delprim* or indole in *Aventicum*, whereas fungal infection and herbivory strongly increase emissions rates and blend complexity. Caterpillar infestation induced characteristic herbivore-associated blends dominated by green leaf volatiles (GLVs; e.g., (*E*)*-*2-hexenal), monoterpenes (e.g., β-myrcene), sesquiterpenes (e.g., (*E*)-β-farnesene), homoterpenes (e.g., (*E*)-4,8-dimethyl-nonatriene; DMNT) and aromatic compounds (e.g., indole) (Fig. 1B; Fig. S1; Tables S1 and S2). Induced blends differed between maize varieties; for example, caterpillar infestation elicited the detectable emission of (*E*,*E*)-4,8,12-trimethyl-1,3,7,11-tridecatetraene (TMTT) in *Delprim* but not in *Aventicum* (Fig. 1B; Tables S1 and S2). Volatile profiles induced by *S. frugiperda* and *S. exigua* showed partial separation in the NMDS ordination. Although the induced blends were qualitatively similar, *S. frugiperda* generally elicited lower emission rates of several compounds, including indole, with these differences being more pronounced in *Delprim* than in *Aventicum* (Fig. 1B; Tables S1 and S2). In contrast, *C. graminicola* infection induced a subset of compounds also detected in caterpillar-infested plants, but typically at lower emission rates. Fungal infection was further distinguished by the release of fatty aldehydes (e.g., octanal), which were not detected in caterpillar-induced profiles, as well as a small number of infection-associated terpenes such as D-limonene (Fig. 1B; Fig. S1; Tables S1 and S2). Consistent with these patterns, Random Forest (RF) classification based on induced maize volatile profiles showed clear discrimination among undamaged, fungus-infected, and caterpillar-infested plants and indicated partial discrimination between the two caterpillar species (Fig. 1C).

#### 1.2) MSS module

An MSS module comprising twelve channels coated with different adsorbent materials (Table S3) was used to measure maize volatile emissions under laboratory conditions, using the same dynamic headspace sampling setup as for GC–MS (Fig. S2). MSS measurements generated distinct electrical response patterns across channels (Fig. 2A). Using MSS responses extracted from a 10-s sampling period, NMDS revealed clear separation among undamaged, fungus–infected, and caterpillar-damaged plants for both maize varieties (Fig. 2B; PERMANOVA: *Delprim*, F₃,₅₉ = 90.086, p = 0.001; *Aventicum*, F₃,₆₁ = 66.422, p = 0.001), with only limited overlap observed between control and fungus–infected plants in *Delprim* and between *S. exigua*– and *S. frugiperda*–infested plants in *Aventicum*. Similarly, RF classification correctly assigned the vast majority of samples to their respective treatments: MSS data enabled clear discrimination not only among undamaged, fungus-infected, and caterpillar-damaged plants, but also between plants attacked by the two closely related caterpillar species.

**Fig. 2.**
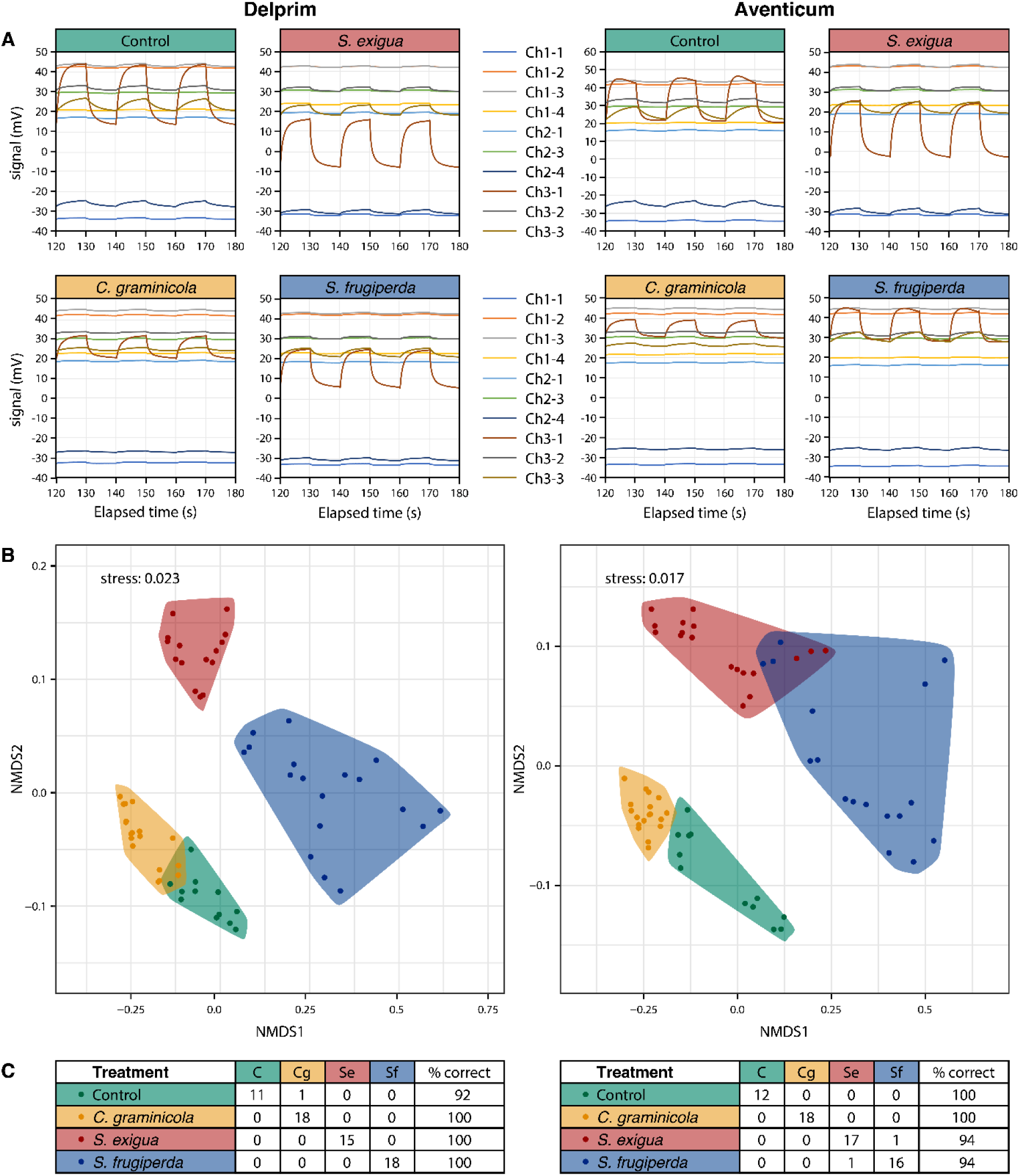
Distinction among maize volatile profiles using the MSS module under laboratory conditions. Volatile emissions were collected under laboratory conditions using the same headspace sampling system as for GC–MS analyses. Undamaged maize plants (green) of the varieties *Delprim* and *Aventicum* were compared with plants infested by a fungal pathogen (*Colletotrichum graminicola*; yellow) or by caterpillars (*Spodoptera exigua*, red; *Spodoptera frugiperda*, blue). **(A)** Sensor signals recorded over three consecutive sampling–purging cycles from ten independent MSS channels with membranes coated with different sensing materials; data from a single 10-s sampling cycle were used for analysis. **(B)** NMDS ordination illustrating differences in sensor response patterns among treatments.**(C)** Confusion matrix summarizing the performance of a Random Forest classification model in assigning measurements to the correct treatment; prediction accuracy was estimated using the model’s internal out-of-bag samples.

#### 1.3) PTR-TOF (Vocus S)

Laboratory measurements of maize volatile emissions were performed with PTR-TOF under laboratory conditions, using the same dynamic headspace sampling setup as for GC–MS (Fig. S2). Volatile compounds previously identified by GC–MS, along with additional ions, were quantified from spectra acquired during 60-s measurements and analyzed as time-averaged concentrations (Fig. 3A; Tables S4 and S5).

**Fig. 3.**
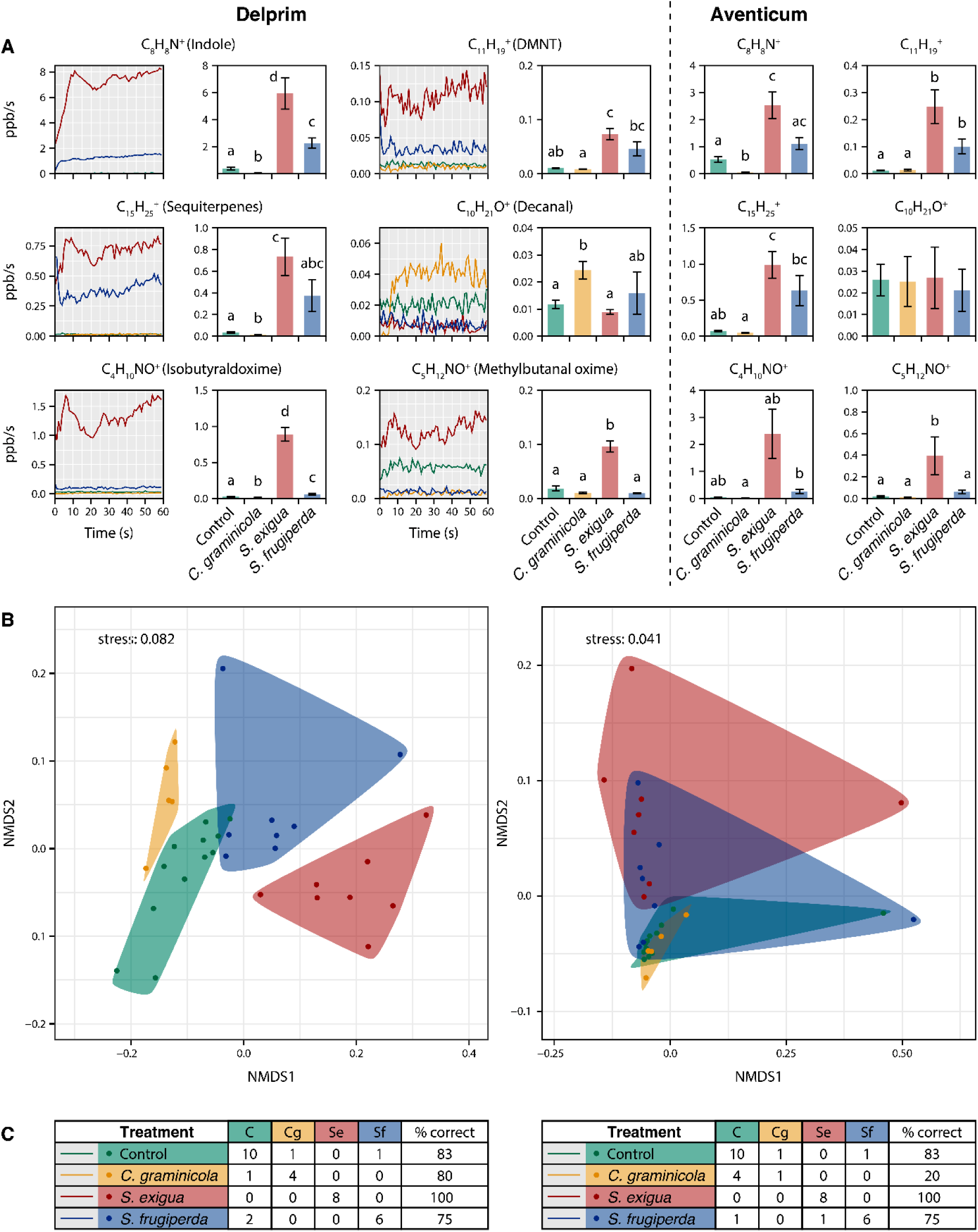
Distinction among maize volatile profiles using PTR-TOF under laboratory conditions. Volatile emissions were measured under laboratory conditions using the same headspace sampling system as for GC–MS analyses. Undamaged maize plants (green) of the varieties *Delprim* and *Aventicum* were compared with plants infested by a fungal pathogen (*Colletotrichum graminicola*; yellow) or by caterpillars (*Spodoptera exigua*, red; *Spodoptera frugiperda*, blue). (**A**) Time series of concentrations (ppb) of selected volatile compounds measured by PTR-TOF over 60-s measurements (1 spectrum s⁻¹), together with bar plots showing mean ± s.e. of concentrations averaged over the 60-s measurement. Different letters indicate significant pairwise differences after FDR correction; full results in Supplementary Information. **(B)** NMDS ordination illustrating differences in volatile profiles among treatments. **(C)** Confusion matrix summarizing the performance of a Random Forest classification model in assigning measurements to the correct treatment; prediction accuracy was estimated using the model’s internal out-of-bag samples.

Several targeted ions showed significant increases in caterpillar-infested plants and corresponded to GC–MS–identified marker compounds, including indole (C₈H₈N⁺) and sesquiterpenes (C₁₅H₂₅⁺), which together represented the most abundant signals, as well as other herbivory-associated compounds such as DMNT (C₁₁H₁₉⁺) and monoterpenes (C₁₀H₁₇⁺) (Fig. 3A; Tables S4 and S5). In addition, PTR-TOF detected several ions associated with caterpillar herbivory that were not observed in the GC–MS analyses, including ions consistent with protonated aliphatic aldoximes (e.g., isobutyraldoxime, C₄H₁₀NO⁺; methylbutanal oxime, C₅H₁₂NO⁺) and other nitrogen-containing compounds such as isobutyronitrile (C_4_H_8_N^+^). These signals were particularly associated with *S. exigua* damage in both maize varieties, although responses were more variable in *Aventicum* (Fig. 3A; Tables S4 and S5). In contrast, ions corresponding to fatty aldehydes characteristic of *C. graminicola* infection in the GC–MS analyses showed little differentiation among treatments in the PTR-TOF data, with the exception of decanal (C₁₀H₂₁O⁺), which was more elevated in infected *Delprim* plants (Fig. 3A; Tables S4 and S5).

For the *Delprim* variety, NMDS revealed clear separation among volatile profiles of all treatments (Fig. 3B; PERMANOVA, F₃,₂₉ = 5.633, p = 0.001), with limited overlap observed between control plants and *S. frugiperda*–infested plants. Accordingly, RF classification distinguished treatments reasonably well, with misclassifications occurring primarily between these two groups (Fig. 3C). For *Aventicum*, NMDS showed more extensive overlap among treatments, although differences in volatile profiles remained detectable (PERMANOVA, F₃,₂₉ = 2.539, p = 0.028; Fig. 3B), most notably between *S. exigua*–infested plants and undamaged or *C. graminicola*–infected plants. Consistent with this pattern, RF classification separated *S. exigua*–infested plants from the other treatments and more generally distinguished caterpillar-infested plants from other treatments but did not discriminate at all *C. graminicola*–infected plants from undamaged controls (Fig. 3C).

### 2) Open-air measurements

Following the encouraging performance of the MSS module and PTR-TOF in laboratory assays, volatile emissions from maize plants (var. *Delprim*) were measured outdoors under open-air conditions without headspace enclosure. Greenhouse-grown plants previously exposed to 24 h of *S. exigua* feeding in the laboratory and subsequently cleared of insects, as well as undamaged controls, were sampled individually immediately outside a campus building. Plants were thus exposed to ambient airflow and environmental conditions, while the surroundings remained relatively simple and non-natural (Fig. S3). Measurements were conducted over three days to incorporate some variability.

#### 2.1) MSS module

MSS measurements were performed at a distance of 1–2 cm from the leaves (Fig. S3). Principal component analysis (PCA) of MSS responses extracted from a 10-s sampling period showed that the dominant sources of variation were associated with sampling day and environmental factors such as temperature and humidity, with no clear separation between damaged and undamaged plants (Fig. S4). Consistent with this result, machine learning models trained on the MSS data did not perform better than random classification. These findings indicate that, under outdoor open-air conditions, MSS signals were strongly influenced by environmental variability and did not contain sufficient treatment-specific information to reliably discriminate plant damage status.

#### 2.2) PTR-TOF (Vocus S)

PTR-TOF measurements were conducted for 60 s at distances of 1–2 cm and 4–5 cm from the leaves under open-air conditions (Fig. 5; Fig. S3), focusing on ions previously identified as herbivore-induced in laboratory assays. PCA of concentrations averaged over the 60-s measurement period indicated that overall variability was primarily associated with sampling day, while plant treatment represented a clear source of variation; temperature and humidity contributed comparatively little (Fig. S5). Visually assessed leaf damage varied substantially across sampling days (median damaged surface: 5%, 15%, and 30%) and ranged thus from minor to relatively severe (Toepfer et al., 2021; Kasoma et al., 2020; Overton et al., 2021). However, concentrations of key herbivore-induced volatiles did not scale with damage extent (Fig. S6), consistent with previous findings that even limited herbivory can already elicit strong volatile responses in maize (Gouinguené et al., 2003).

Several ions were significantly elevated in damaged plants at both sampling distances, including indole (C₈H₈N⁺), sesquiterpenes (C₁₅H₂₅⁺ and fragment ions), isobutyraldoxime (C₄H₁₀NO⁺), DMNT (C₁₁H₁₉⁺), and jasmone (C₁₁H₁₇O⁺) (Fig. 4A; Table S6). Others, such as monoterpenes (C₁₀H₁₇⁺) and TMTT (C₁₆H₂₇⁺), showed significant differences only at the shorter sampling distance (Table S6). Concentrations detected during open-air measurements were markedly lower than those measured under laboratory enclosed headspace conditions, typically reaching low ppt levels (Table S6). Median indole (C₈H₈N⁺) concentrations decreased by more than 200-fold from laboratory measurements to open-air sampling at 1–2 cm and by nearly 300-fold at 4–5 cm, while median sesquiterpene (C₁₅H₂₅⁺) concentrations declined by 26-fold at 1–2 cm and 50-fold at 4–5 cm. In comparison, the additional attenuation between 1–2 cm and 4–5 cm was modest (approximately 1.3–2-fold) (Table S7).

**Fig. 4.**
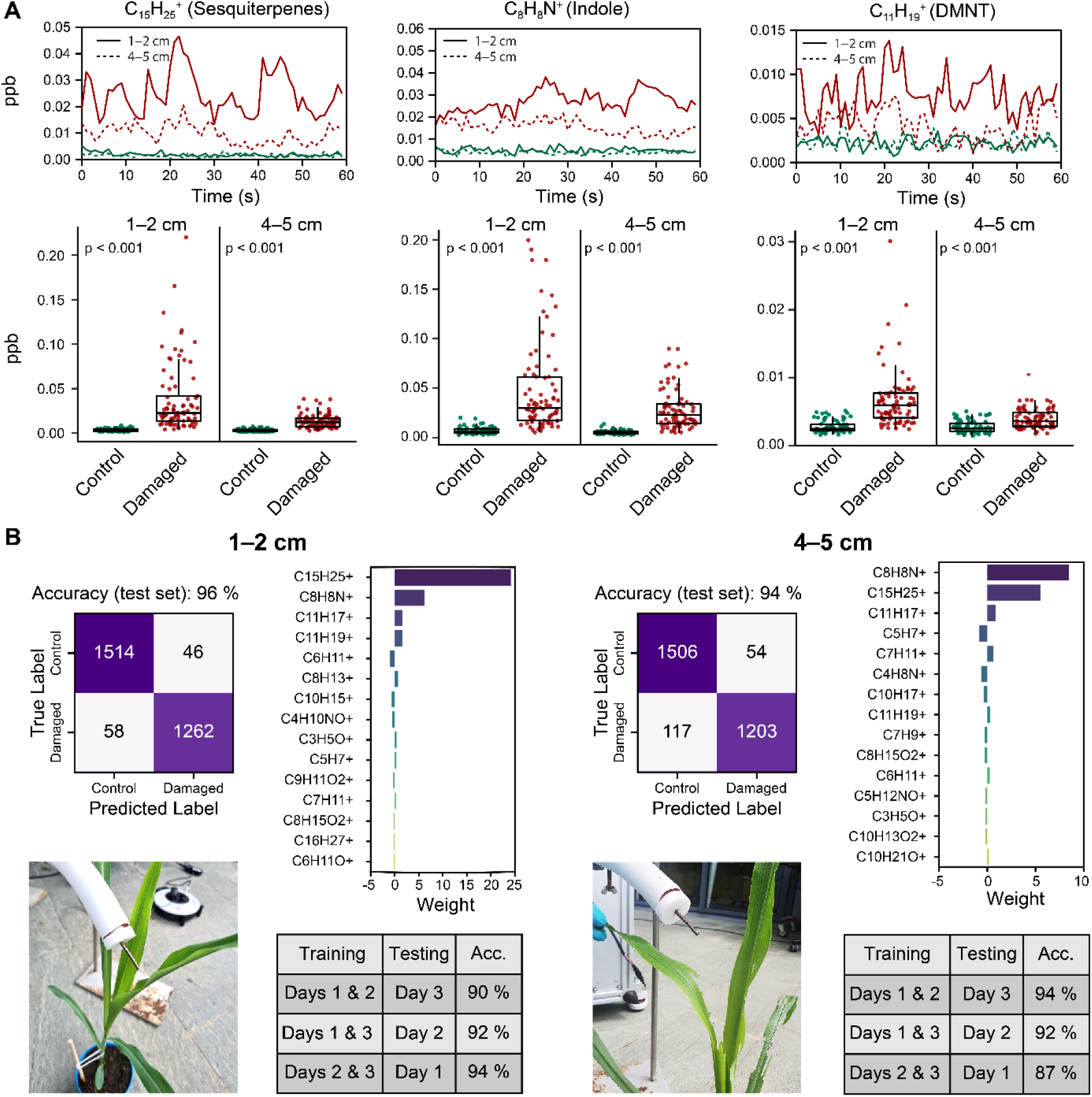
Distinction between volatile emissions from undamaged and damaged maize plants under outdoor, open-air sampling, conditions using PTR-TOF. Volatile emissions were measured under open-air conditions from maize plants (variety *Delprim*) using PTR-TOF. Control (undamaged) plants (green) were compared with plants previously damaged in the laboratory for 24 h by *Spodoptera exigua* caterpillars; removed prior to measurements. In total, 77 control and 81 damaged plants were measured over three days. For each plant, 60-s measurements were performed either close to the leaves (1–2 cm) or at a greater distance (4–5 cm). **(A)** Time series of concentrations (ppb) of selected volatile compounds recorded at 1-s resolution over 60 s, together with boxplots showing concentrations averaged over the 60-s measurement (individual data points shown). P-values for treatment comparisons are shown (Wilcoxon rank-sum tests; full results in Supplementary Information). **(B)** Performance of classification models based on Lasso-regularized logistic regression used to distinguish undamaged from damaged plants using volatile profiles. The confusion matrix summarizes predictions on an independent test set (30% of plants), after model training on the remaining 70% of plants; overall classification accuracy is indicated. For model training and testing, each 60-s measurement was divided into 60 consecutive 1-s observations, and all observations from a given plant were assigned to the same training or test set. The importance plot ranks the 15 most influential volatile compounds based on their model coefficients, where larger absolute values indicate a stronger contribution to distinguishing damaged from undamaged plants. The table reports predictive accuracy for models trained on data from two sampling days and tested on data from a third, independent day.

To evaluate the potential for damage detection under open-air conditions, machine learning models were trained to predict plant damage status from PTR-TOF measurements. Subsampling each 60-s measurement into shorter intervals (1, 5, 15, 30 s) showed that even single-second measurements contained sufficient information for accurate classification (Table S8). Lasso-regularized logistic regression yielded the best performance and was therefore used for further tuning. Models based on 1-s intervals correctly predicted damaged and undamaged plants in independent test datasets with accuracies of 96% at 1–2 cm and 94% at 4–5 cm (Fig. 4B).

Analysis of model coefficients revealed strong sparsity in predicting features, with only a small number of ions contributing substantially to damage prediction and others either weakly weighted or shrunk to zero by the regularization. At the shorter sampling distance (1–2 cm), sesquiterpenes (C₁₅H₂₅⁺) were the dominant predictors, while indole (C₈H₈N⁺) also contributed substantially (Fig. 4B). At the larger sampling distance (4–5 cm), the relative importance of these predictors shifted, with indole emerging as the strongest contributor, followed by sesquiterpenes (Fig. 4B).

To assess robustness to day-to-day variation in PTR-TOF signal structure, models were trained on data from two sampling days and applied to predict plant damage status on a third, previously unseen day. These models achieved classification accuracies of ≥86% (Fig. 4B), indicating that, despite clear day-to-day variability in measurements (Fig. S5), treatment-associated signals were sufficiently stable to enable accurate cross-day damage classification.

#### 2.3) Field measurements with ultra-portable TOF (Vocus C)

As real-time mass spectrometry proved effective for the rapid open-air detection of volatile markers associated with herbivory damage, we next assessed the feasibility of this approach under increasingly realistic conditions by performing in situ measurements on field-grown plants (var. *Delprim*). As logistical and infrastructure requirements of deploying standard laboratory instrument in real world field conditions are unrealistic in an off grid scenario, we used a recently developed ultra-portable TOF (Vocus C). It weighs around 30kg and consumes only 250W of power, overcoming the main practical limitations of making field measurements. The instrument was operated with benzene cation ionization which is a more selective chemical ionization approach than PTR based ionization.

We first tested the instrument on greenhouse-grown maize plants subjected to simulated herbivory in direct comparison with PTR-TOF (Fig. S7) in the same open-air setup. Under these conditions, the ultra-portable TOF reliably detected key herbivory-associated volatiles, including indole and sesquiterpenes, with performance comparable to PTR-TOF (Fig. S7).

Field measurements were then conducted on maize plants grown in an agricultural field plot in Mathod, Switzerland (Fig. S8). Plants subjected to simulated herbivory were individually sampled for 60-s (Fig. 5A). Raw ion count data, binned by nominal m/z value across the full mass spectrum and averaged over the 60-s measurement period, were used for analyses. To assess whether measurements acquired under field conditions could discriminate between damaged and undamaged plants, we applied a Lasso-regularized logistic regression model. Across repeated cross-validation, the model achieved a mean classification accuracy of 69.0 ± 2.6 % (95% CI), indicating stable performance across resampling (Fig. 5B). Although classification accuracy was moderate, these results indicate that real-time mass spectrometry captured damage-related information from plant volatile profiles under realistic field conditions.

**Fig. 5.**
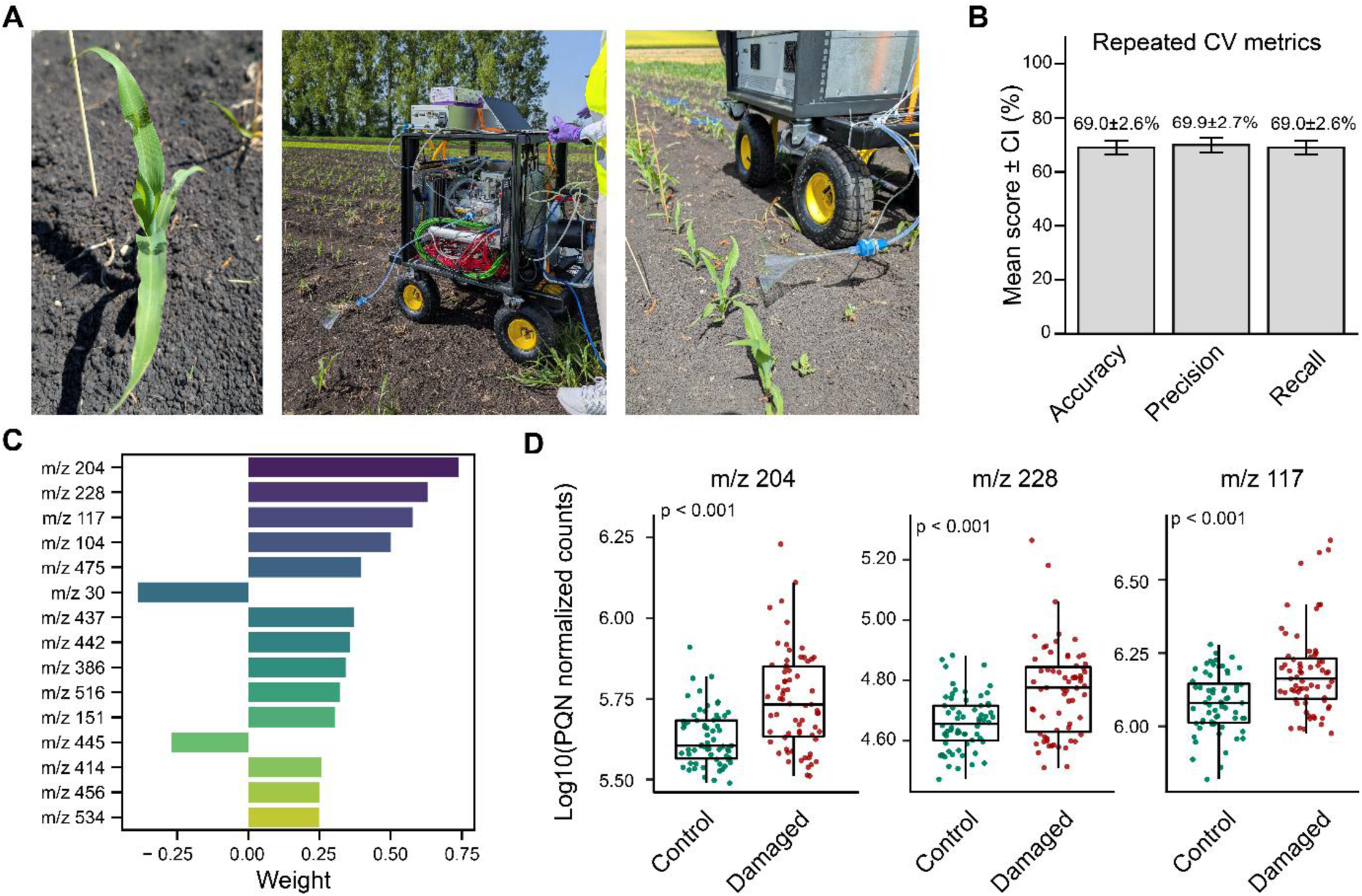
Preliminary field validation of maize volatile profiling for detecting herbivory damage using ultra-portable TOF. Measurements were performed on maize plants under field conditions using a portable instrument. Simulated herbivory consisted of mechanical leaf damage combined with the application of caterpillar regurgitant. **(A)** Photographic overview of the experimental setup, showing (i) a maize plant subjected to simulated herbivory in the field, (ii) the instrument mounted on a mobile cart, and (iii) the sampling configuration **(B)** Classification performance for distinguishing damaged and undamaged plants using Lasso-regularized logistic regression. Analyses were performed in a non-targeted manner using ion intensities at each nominal m/z value, averaged over 60-s measurement period, probabilistic quotient normalized (PQN) and log₁₀-transformed prior to modeling. Model performance was evaluated using repeated cross-validation. Bars show mean performance metrics across repeated cross-validation, with error bars indicating 95% confidence intervals. Accuracy reflects the overall proportion of correctly classified plants, while precision and recall describe the rates of false-positive and false-negative classifications, respectively. **(C)** Feature importance ranking derived from the same cross-validation procedure. Bars show mean model coefficients across cross-validation runs, reflecting the relative contribution of individual m/z features to discrimination between damaged and undamaged plants. Only the 15 most influential features are shown. **(D)** Boxplots showing log-transformed and normalized signal intensities for the three most informative m/z features identified in (C). Individual plant data points are shown; p-values are from Wilcoxon rank-sum tests (full results in Supplementary Information).

Analysis of feature importance, based on mean model coefficients across cross-validation splits, identified multiple m/z features contributing to discrimination between damaged and undamaged plants (Fig. 5C), indicating a more complex feature structure than observed in the less realistic open-air experiment. The three most important features corresponded to sesquiterpenes (m/z 204), a benzene-adduct of DMNT (m/z 228), and indole (m/z 117), all of which exhibited significantly higher signal intensities in damaged plants (Fig. 5D). Several additional m/z features also showed non-zero coefficients but were not assigned here. The prominence of indole- and sesquiterpene-related signals among the top predictors mirrors our open-air PTR-TOF analyses, indicating that these key herbivory markers remained detectable and informative under realistic field conditions. Together, these findings support the feasibility of real-time mass spectrometry for open-air monitoring of herbivore damage in agricultural settings.

## Discussion

Plants rapidly emit specific volatile compounds in response to biotic threats, a well-documented process central to plant defense and ecological signaling (Baldwin, 2010; Heil, 2014; Schuman, 2023; Turlings and Erb, 2018). This has motivated the idea of harnessing plant odors as early indicators of biotic stress in crops (Haque et al., 2026; Brilli et al., 2019; Pickett & Khan, 2016). In this study, we evaluated the feasibility of such an approach in maize using currently available sensing technologies. We adopted a stepwise experimental strategy, progressing from laboratory assays with enclosed plants to open-air measurements in a simple outdoor setting and, finally, to an initial field trial. This allowed us to assess whether induced plant volatile signatures remain detectable and informative as conditions shift from controlled to progressively more realistic settings.

### Volatile signatures of biotic attacks in maize

Laboratory GC-MS analyses confirmed that maize plants emit distinct volatile signatures in response to herbivory and fungal infection. Caterpillar feeding induced the release of multiple terpenes, GLVs, and aromatic compounds, whereas undamaged plants emitted few volatiles at low abundance, consistent with prior studies (D’Alessandro and Turlings, 2006; De Lange et al., 2020). The two *Spodoptera* species elicited quantitatively distinct responses, as previously described (De Lange et al., 2020), illustrating that volatile signatures can be sufficiently nuanced to potentially distinguish between closely related pests. While such discrimination may not be essential for practical pest detection if management strategies are similar, it highlights the rich information content of herbivore-induced volatile blends. Fungal infection elicited a subset of compounds also observed during herbivory, but additionally induced characteristic fatty aldehydes and monoterpenes, consistent with previous reports for *Colletotrichum* sp. in other crops (Quintana-Rodriguez et al., 2015), although volatile emission responses of maize to infection by Colletotrichum pathogens remain poorly characterized and largely undocumented.

Importantly, volatile responses varied between maize varieties, with one variety showing clearer separation among treatment-specific volatile profiles than the other. Such intraspecific variation is well documented in maize and other crops (Gouinguené et al., 2001; Degen et al., 2004; Hare, 2011; Martín-Cacheda et al., 2023), and our results further suggest that the performance of odor-based monitoring tools may depend on plant genotype. This implies that detection approaches should be evaluated across multiple varieties to assess their robustness and general applicability. At the same time, this variability also offers opportunities to enhance detection by selecting or prioritizing cultivars that emit stronger or more distinct stress-induced volatile signals.

### Odor sensing under laboratory conditions with enclosed headspace sampling

Under enclosed headspace sampling, The MSS module performed very well in laboratory assays, reliably distinguishing healthy maize plants from infested and infected plants, and discriminating among different attackers. This finding adds to a few studies demonstrating successful use of portable odor sensors for discriminating different types of biotic stress through plant induced volatiles, which have likewise relied on enclosed sampling conditions, either static or dynamic (Laothawornkitkul et al., 2008; Zhou and Wang, 2011; Ghaffari et al., 2012; Sun et al., 2019; Cui et al., 2021).

PTR-TOF also distinguished among treatment-specific volatile signatures in laboratory assays, with particularly strong performance for herbivory. In addition to detecting the major compounds identified by GC-MS, it provided complementary information on the herbivore-induced volatile blend by capturing likely aldoximes and other nitrogen-containing compounds that were not detected by GC-MS. Such compounds have been reported as components of herbivore-induced volatile emissions in several plant species (Sørensen et al., 2018) and, occasionally, in maize (Takabayashi et al., 1995; Turlings et al., 1998a). These results underscore that the full volatile response of plants can be richer than is typically captured by GC-MS alone, and that minor compounds may contribute to valuable chemical information that can improve the accuracy and specificity of odor-based plant stress detection tools.

At the same time, PTR-TOF also has inherent limitations. Its performance in identifying fungal infection was less consistent and varied between maize varieties. Fatty aldehydes, which were characteristic markers of fungal infection in GC-MS analyses, were poorly detected, likely because their parent ions undergo extensive fragmentation during PTR ionization, generating overlapping masses that reduce chemical specificity (Buhr et al., 2002; Kari et al., 2018; Ernle et al., 2023). Alternative ionization strategies, for instance adduct-ionization (Riva et al., 2024), may therefore be required to improve detection of such functionalized volatiles.

### Odor sensing in open-air conditions

Under open-air conditions, volatile concentrations were strongly diluted relative to laboratory assays, as expected. PTR-TOF measurements indicated that herbivore-induced compounds occurred at low ppt levels, more than two orders of magnitude lower than in enclosed headspace experiments. Under these conditions, the MSS module did not capture treatment-specific volatile signatures of herbivory damage. This lack of discrimination likely reflects a combination of strong dilution and the influence of fluctuating environmental factors, such as humidity and temperature, on sensor performance. These observations are consistent with previous reports identifying sensitivity constraints and environmental dependence as major challenges for deploying portable gas sensors for plant odor detection under field conditions (Cui et al., 2018; Ivaskovic et al., 2021; MacDougall et al., 2022; Zheng and Zhang, 2022; Chen et al., 2024, p. 202). In addition, as an equilibrium-based sensor, MSS may be particularly susceptible to rapid concentration fluctuations because the receptor layer requires time to equilibrate with the surrounding air, reducing signal strength under dynamic open-air conditions. Although the MSS module can reach ppt-level detection limits for some compounds (Minami et al., 2018), the time required to accumulate sufficient signal at very low concentrations currently exceeds one minute (Minami et al., 2021, 2023), limiting its suitability for mobile, continuous monitoring in crop environments. Nonetheless, the compact size, ease of use, and strong performance under more controlled sampling conditions at detecting induced plant volatile signatures suggest potential for other on-site applications where volatile concentrations are higher or sampling can be stabilized, such as inspections in containers of imported plant material (Favaro et al., 2024; Karimi and Gross, 2024; Asiri et al., 2024).

PTR-TOF detected several herbivore-induced volatiles under open-air conditions, even at low ppt concentrations. In this simple experimental setup, data collected at one-second intervals contained sufficient information to distinguish herbivory-damaged from undamaged plants with very high accuracy, with indole and sesquiterpenes showing to be particularly informative markers. Importantly, this does not imply that other volatiles were uninformative; rather, these two compounds alone were sufficient to achieve near-perfect discrimination. This strong predictive performance, driven largely by only two markers, likely reflects the experimental and biological simplicity of the system. Plants were grown under optimal and constant greenhouse conditions and assessed in a simple outdoor setting that does not capture the complexity of natural crop environments. Under these conditions, undamaged plants emitted little to no detectable volatiles, whereas damaged plants produced clear signals, effectively reducing discrimination to the presence or absence of a small number of dominant characteristic compounds. Moreover, plants were measured individually, without interference from neighboring plants, making odor attribution straightforward. In field settings, by contrast, plants are exposed to multiple and variable biotic and abiotic stresses that could shape volatile responses (Holopainen and Gershenzon, 2010; Turlings and Erb, 2018; Schuman, 2023), and their emissions could overlap with odor plumes from different plants and other sources. Reliable detection under such conditions will therefore likely require integrating information across a broader set of volatiles and applying full fingerprint-based pattern-recognition approaches, as implemented in our field assay, rather than relying on a few dominant markers alone. Although our open-air setup does not capture this complexity, the results demonstrate that real-time mass spectrometry can rapidly resolve low-abundance plant odor signals under moderately variable environmental conditions (e.g., day-to-day changes in humidity and temperature), providing a foundation for extending measurements to more complex and heterogeneous field scenarios.

### Odor sensing under field conditions

To evaluate odor-based detection under realistic agricultural conditions, we conducted an initial field trial. Although limited in scope (a single sampling day and one field plot), it supports the feasibility of real-time detection of herbivory damage through induced volatiles under true field conditions. Using a portable TOF, damaged plants could be distinguished from undamaged plants with an accuracy of almost 70%, a level of performance that is particularly encouraging given that neither the instrumentation nor the sampling strategy was specifically optimized for field-based plant odor detection at this stage.

As expected, compared with the less realistic open-air experiment, discrimination under field conditions was characterized by greater signal overlap and a more complex structure of discriminatory features. Despite this increased complexity, the most influential discriminating signals again corresponded to indole, sesquiterpenes, and very likely DMNT, indicating that these key herbivory-associated volatiles remain detectable and informative even under realistic field conditions.

However, extending odor-based detection using real-time mass spectrometry toward practical deployment will require evaluating the robustness and specificity of volatile-based diagnostics under natural sources of variability. A central challenge will be determining whether different plant stresses can be reliably distinguished when they co-occur simultaneously and potentially generate overlapping odor plumes. Addressing this complexity will require testing across a range of plant developmental stages, environmental conditions, and past and present stress scenarios. This, in turn, will necessitate extensive spatial and temporal sampling across heterogeneous agricultural environments and the development of machine-learning–based pattern-recognition approaches to integrate high-dimensional, dynamic odor data and enable automated, real-time diagnostic interpretation.

Practical implementation is further constrained by airflow conditions that are more difficult to control in field environments than in simplified outdoor experiments. In the latter, wind direction was relatively stable, partly constrained by surrounding buildings, allowing sampling inlets to be positioned consistently downwind of individual plants, while wind speeds remained low (< 3 m s⁻¹). Under field conditions, wind speeds were similarly low (< 3 m s⁻¹), but wind direction was inherently more variable, which can dilute or redirect emissions and increase contributions from neighboring plants, complicating odor attribution. These limitations are expected to become more pronounced under stronger wind conditions, which would further disrupt plume structure and sampling geometry. These constraints can be overcome with refined sampling strategies, such as partial airflow shielding, adaptive inlet positioning, or incorporation of wind direction information into odor source-mapping strategies (Loubet et al., 2022; Sarkar et al., 2020; Petersen et al., 2025).

Despite these limitations and the clear need to repeat and expand field experiments to ensure generalizability, the results are encouraging, highlighting the potential of odor-based detection approach using real-time chemical ionization mass spectrometry. Signal strength is likely to increase under more realistic infestation scenarios. Notably, our experimental layout likely represents a particularly challenging configuration, as very small sets of damaged and undamaged plants were positioned adjacent to each other’s, requiring fine-scale detection. In contrast, natural infestations typically occur in spatial clusters rather than as isolated plants (Karimzadeh and Sciarretta, 2022; Bhargav et al., 2025; Zanzana et al., 2025), creating larger aggregated sources of herbivore-induced volatiles rather which should generate stronger signals and make detection less sensitive to small variations in wind direction. In addition, continuous herbivory is expected to produce stronger and more sustained volatile emissions than the temporally limited simulated damage applied here (Turlings et al., 1998a).

### Outlook

Overall, this study provides an initial proof of concept for odor-based detection of agricultural pests based on induced plant volatile emissions. Real-time mass spectrometry enabled sensitive and temporally resolved detection of herbivory-associated signatures under open-air conditions, and preliminary field data further support the feasibility of in situ plant stress monitoring. At the same time, these findings require confirmation through more extensive field validation before practical deployment can be considered. Moreover, at present, CI-TOF instruments remain costly, limiting their direct deployment in agricultural settings. Nevertheless, even if routine field use remains impractical in the near term, their ability to identify and quantify informative marker volatiles under realistic conditions provides valuable guidance for the development of next-generation portable sensors. Because many crop species emit overlapping volatile profiles in response to herbivory (Turlings and Wäckers, 2004), insights gained here should be transferable beyond maize. In a broader context, odor-based monitoring could complement existing precision agriculture tools, such as robotic weeders and selective sprayers, that aim to reduce agrochemical inputs (Anne et al., 2024; Lochan et al., 2024; Ye et al., 2025). It may also support more targeted and cost-effective deployment of biological control agents (Cusumano et al., 2020; Haque et al., 2026), including microbial insecticides and entomopathogenic nematodes (Gardner and Fuxa, 1980; Fallet et al., 2022). In conclusion, while substantial challenges remain, our findings highlight the potential of plant volatile sensing as a component of sustainable crop protection and motivate further interdisciplinary research to assess its robustness, applicability and scalability under agricultural conditions.

## Methods

### Plants and growth conditions for laboratory and open-air assays

Maize plants, *Zea mays var.* Delprim and *var.* Aventicum (seeds supplied by Delley Semences et Plantes SA, CH), were grown in plastic pots (4 cm diameter × 10 cm height for laboratory experiments ; 10 cm diameter x 8.6 cm height). Pots were filled with commercial potting soil (Ricoter, Erdaufbereitung AG, CH) and grown under greenhouse conditions (26 ± 2°C, 60 ± 5% relative humidity). Natural sunlight was supplemented with LED lamps (luminous flux 8,800 lm), providing a 16 h photoperiod. Plants were watered every other day and fertilized once per week with an 8–8–6 NPK fertilizer (Maag Garden, Switzerland). Plants used for laboratory measurements were 10-12 days old and had 2-3 leaves. For the open-air measurements, plants were 20-25 days old and had 4-5 developed leaves.

### Pathogen, insects and regurgitant collection

#### Fungus

The fungal pathogen *Colletotrichum graminicola* (Ces.) G.W., which causes maize anthracnose leaf blight disease, was used to inoculate maize plants for laboratory measurements. The inoculation was done following (Balmer et al., 2013). Fungal colony (M1.001, obtained from Lisa Vaillancourt, University of Kentucky, Department of Plant Pathology, Lexington, KY) was maintained at 25°C on potato dextrose agar (Difco PDA, Becton Dickinson) under continuous light (70 µE m^−2^ sec^−1^). Three-week-old plates were used to harvest spores for infection. Twenty µL of a *C. graminicola* conidia suspension (6 × 10^5^ spores mL^−1^ sterile water with 0.01% Silwet L-77, Lehle Seeds) were applied and spread over the leaves using a syringe, four days prior to odor collections. After inoculation, infected plants were kept under room conditions for 16h, without direct sunlight and in an open plastic box (50 × 30 × 35 cm) containing vaporized water used as a humid chamber (min 75% relative humidity, 23 ± 2°C), before transferring them back to an isolated place in the greenhouse. Fungus-inoculated plants were further monitored after odor measurements, as signs of a successful anthracnose leaf blight disease were hardly detectable at this stage. Plants that had not developed any disease symptoms on day 15 (i.e., a week after inoculation) were excluded from further analyses.

#### Caterpillars

In-house colonies of the beet armyworm *Spodoptera exigua* and the fall armyworm *Spodoptera frugiperda* (OFEV permits A192632-07 and A192558-02) caterpillars were reared on a wheat germ-based artificial diet (Frontier Scientific Services, Newark, USA) in transparent plastic boxes under laboratory conditions (temperature 30 ± 5°C, relative humidity 60–80%, and 16 h photoperiod) (Arce et al., 2021). For laboratory measurements, second instar *S. exigua* and *S. frugiperda* larvae were used to damage the plants. For open-air measurement, late third instar *S. exigua* were used. In all cases, volatile sampling was performed 24 h after the onset of infestation.

#### Regurgitant collection

Caterpillar regurgitant, used for simulating herbivory, was collected as previously described (Arce et al., 2021). Briefly, third- to fourth-instar *S. frugiperda* larvae were fed on maize leaves (Delprim) for 24 h. Larvae were gently squeezed in the head region using featherweight entomological forceps to induce regurgitant release, which was collected into Eppendorf tubes and stored at −80 °C until use.

### Laboratory assays using GC-MS, MSS and PTR-TOF

Volatile emissions from maize plants (Delprim and Aventicum) attacked by herbivores or pathogens were measured using dynamic headspace sampling, where each plant is enclosed in a one-port glass bottle as described by (Turlings et al., 2004). Charcoal-filtered, humidified air entered the bottle at 0.7 L min^−1^. This was combined with three analytical techniques: gas chromatography–mass spectrometry (GC-MS), membrane-type surface stress sensors (MSS), and proton-transfer-reaction time-of-flight (PTR-TOF) mass spectrometry (Figs. S1 and S2).

#### GC-MS

Volatiles were collected from maize plants infested with either six *S. exigua* (n = 7–8) or four *S. frugiperda* (n = 7) second-instar larvae per plant. The different larval numbers were chosen to achieve comparable feeding damage, as *S. frugiperda* feeds more intensively on Poaceae (Kenis et al., 2023). Sampling was conducted 24 h after infestation. For fungal treatments, plants inoculated with *C. graminicola* were sampled seven days post-inoculation (n = 8–9). Control plants received no treatment (n = 8). Air was pulled through a 25 mg of Porapack Q 80/100 mesh Hayesep-Q adsorbent filter (Ohio Valley Specialist Company, arietta, USA) at 0.4 L min^−1^. Volatiles were collected for 2 h. Filters were eluted with 150 µL dichloromethane (Honeywell, Riedel-de Haën, Germany), and 10 µL of internal standard (n-octane and n-nonyl acetate, 20 ng µL⁻¹ each) were added. Samples were stored at −80 °C until analysis. Analyses were performed using an Agilent 7890B gas chromatograph coupled to an Agilent 5977B mass spectrometer operating in total ion current (TIC) mode. A 1.5 µL aliquot was injected in pulsed splitless mode onto an HP-5MS column (30 m × 0.25 mm × 0.25 µm). The oven temperature program was 40 °C for 3 min, increased to 100 °C at 8 °C min⁻¹, then to 230 °C at 5 °C min⁻¹. Helium was used as carrier gas at 1.1 mL min⁻¹. Compounds were identified by comparison with commercial standards and NIST 17 library spectra and quantified by using relative response factors obtained from calibration curves of commercial standards in comparison to nonyl acetate

#### MSS

A nanomechanical Membrane-type Surface stress Sensors standard measurement module (7.5 × 11.5 × 3 cm, developed by MSS Alliance/Forum (Alliance, 2015; Forum, 2017) (Yoshikawa et al., 2011, 2012; Minami et al., 2022) provided by National Institute for Materials Science (NIMS; Tsukuba, Japan) was used to sample volatiles from maize plants infested with three *S. exigua* larvae (n = 15–18), two *S. frugiperda* larvae (n = 18), or inoculated with *C. graminicola* (n = 18). Control plants were untreated (n = 12). The outlet port of the glass bottles was directly connected to the MSS module via Teflon tubing (50 cm length, 1/16″ OD, 1 mm ID), and an empty bottle was used as source of purging air (Fig. S2). The module consists of a compact micro-fabricated array with twelve channels, each containing a 3 µm-thick silicon membrane with a diameter of 300 µm coated with different receptor materials, such as functional silica/titania hybrid nanoparticles or polymers (Table S3), conferring selectivity for gas mixtures (Shiba et al., 2015, 2017; Minami et al., 2022). Molecular adsorption and desorption on the receptor layers deform the membranes, altering the resistance of piezoresistors embedded in four sensing beams. These resistance changes are measured (in mV) using a full Wheatstone bridge with a –1.0 V bridge voltage (Yoshikawa et al., 2011) and recorded at 100 Hz. Measurements are continuous and rely on the alternance of sampling (adsorption phase of the analytes) and purging (acceleration of desorption) phases. Flow rates of both sampling and purging phases were set at 0.030 L min^−1^. Each laboratory measurement consisted of 10 cycles of 10-second sampling and 10-seconds purging. Of the 12 available channels in the module, two showed unstable responses throughout measurements (channels 2.2 and 3.4, Table S3) and were therefore excluded from the analyses. Data from the sampling phase of the 8th cycle were used, extracting one value per second over the final 9 s (141–150 s).

#### PTR-TOF

volatiles were analyzed using PTR-TOF (Vocus S, Tofwerk AG, Thun, Switzerland). Plants were infested with six *S. exigua* (n = 8), four *S. frugiperda* (n = 8), or inoculated with *C. graminicola* (n = 5). Control plants were untreated (n = 12). Air was sampled from the bottles port at 0.1 L min⁻¹ through a 1 m inlet heated to 100 °C to minimize losses. Volatile compounds were ionized by proton transfer from H₃O⁺ reagent ions. Further details about the Vocus S technology can be found elsewhere (Krechmer et al., 2018; Li et al., 2020). The ion–molecule reactor was operated at 1.8 mbar and 100 °C, with front and back voltages of 500 V and 38 V, respectively, and an RF field of 1.3 MHz at 450 V. Mass spectra were recorded at 1 Hz over an m/z range of 20–486.

Raw data were processed using Tofware 3.2.3 (Tofwerk AG, Thun, Switzerland) for mass calibration, peak fitting, and quantification. Targeted ions included parent and characteristic fragment ions of plant volatiles (Tables S4 and S5), selected based on GC-MS data, prior experience, and literature (Yáñez-Serrano et al., 2021). Signal intensities were converted to concentrations (ppb) using instrument responses derived from a standard mixture with known proton-transfer rate constants, as described in Lopez-Hilfiker et al. (2019), ran prior to measurements using a 20 ppb gas mixture (Apel-Riemer Environmental, Miami, USA; using acetaldehyde, acetonitrile, acrylonitrile, acetone, benzene, 1,2,4-trimethylbenzene, toluene, *m*-xylene and methyl ethyl ketone). The estimated uncertainty of this quantification method is approximately 30%.

### Open-air sampling using MSS and PTR-TOF in semi-controlled outdoor conditions

Open-air volatile sampling, without headspace enclosure, was conducted using the same PTR-TOF and MSS systems as described above over three days (1, 2, and 5 July 2024) at the Faculty of Science of the University of Neuchâtel, Switzerland (47°00′00.5″ N, 6°57′00.5″ E) (Fig. S3). Measurements were performed between 09:00 and 14:00 under mostly sunny conditions, with occasional overcast periods. Ambient temperature (19–26 °C), relative humidity (42–60%), and wind speed (1–10 km h⁻¹) were recorded manually every 15–20 min using a portable weather station (TFA Spring Breeze, Dostmann GmbH & Co. KG, Germany). Sampling was carried out immediately outside a university building, where plants were exposed to natural airflow and ambient environmental conditions, while remaining in a semi-sheltered setting: the proximity of the building limited wind exposure to fewer directions. This facilitated positioning the MSS module and PTR-TOF sampling inlets downwind of the plant whenever possible.

Plants were grown in the greenhouse as described above and transferred to a laboratory room one day prior to measurements, where they were maintained under artificial light (200 µmol m⁻² s⁻¹, 16 h : 8 h light/dark, 26 ± 2 °C). Half of the plants were infested with two to three late third-instar *S. exigua* larvae for 24 h, while the remaining plants served as undamaged controls. To prevent larvae from falling off plants during infestation without interfering with feeding behavior, infested plants were partially enclosed in open-ended PET cooking bags secured around the stem. One quarter of control plants were also bagged; this treatment did not induce detectable volatile emissions (Table S9). Bags and larvae were removed 2–3 h before sampling. Feeding damage was assessed visually as the percentage of leaf area removed.

Volatiles were sampled from the space between leaves, 5–15 cm above the whorl, targeting the youngest fully expanded leaf, regardless of whether that leaf itself was damaged (Fig. 4 and Fig. S3). Measurements alternated between control and damaged plants. Each plant was first sampled using PTR-TOF, followed immediately by MSS measurements. PTR-TOF sampling was conducted for 60 s at a distance of 1–2 cm from the leaf and subsequently at 4–5 cm from the same leaf. MSS sampling was performed at 1–2 cm for seven cycles of 10 s sampling followed by 10 s purging. The MSS inlet was reduced to a 16 cm Teflon tube, and technical air was used for purging. The PTR-TOF inlet was fixed on a clamp stand and the maize leaves were delicately held with gloved-fingers to maintain the proper distance, despite the wind. The MSS inlet was similarly maintained at the right distance from leaves (Fig. S3).

On day 1, we measured a total of 52 plants (control n = 25, damaged n = 27), on day 2 a total of 60 plants (control n = 22, damaged n = 28) and on day 3, a total of 56 plants (control n = 30, damaged n = 26). Analyses targeted the same ions as laboratory measurements, restricted to caterpillar-induced compounds (Tables S4 and S5). Quantification was based on a calibration run daily, prior to measurements, as described for the laboratory assay.

### Field measurements with ultra-portable TOF

#### Plot and instrument

The field assay was conducted in an agricultural maize plot in Mathod, Switzerland (46.7493° N, 6.5545° E) (Fig. S8). The experiment took place on 2 May 2025 and was limited to a single day due to weather conditions and instrument availability. Maize plants (var. *Delprim*) grew under standard agronomic conditions. Seeds were sown at the end of March with 25 cm spacing within rows and 50 cm between rows. The field plot measured ∼60 m × 18 m. Two rows were used for the experiment, separated by one untreated row. The first experimental row was located 3 m from the plot edge, and only plants situated at least 3 m from the remaining plot borders. Plants were used at 3 (true) leaves, with the 4^th^ one emerging. Measurements were performed using an ultra-portable TOF instrument (Vocus C, Tofwerk AG, Thun, Switzerland) (Dobrecevich et al., 2026). It weighs approximately 30 kg and has a low power consumption of ∼250 W. The instrument sensitivity and detection limits are comparable to the aforementioned Vocus S PTR-TOF instrument. For this experiment the instrument was powered using the battery of a hybrid car. A catalytic zero air generator (Tofwerk AG, Thun, Switzerland) was used to generate clean air, which was further purified with a VOC trap (VICI XXX) allowing the instrument to be independent of any auxiliary gas supplies. While the compact design of the instrument balances sensitivity, weight and power consumption its mass resolving power is somewhat limited (1200 Th/Th) and is best coupled with a rather selective reagent ion to maintain relatively simple mass spectra. The instrument was operated with benzene cation ionization which is a more selective chemical ionization approach than PTR based ionization (Puttu et al., 2025; Riva et al., 2024; Kim et al. 2016). This ionization method enables detection predominantly through charge transfer processes or benzene adduct formation, depending on the compound properties.

For sampling, air was drawn at 10 L min⁻¹ through unheated PTFE tubing using a glass funnel (10 cm diameter) mounted at the inlet to increase the sampling area and reduce sensitivity to exact inlet positioning.

#### Simulated herbivory

plants were treated in alternating groups of three damaged and five control, undamaged plants across two rows. Herbivory damage was simulated by combining mechanical wounding with the application of *S. frugiperda* regurgitant. On the day before measurements (1 May 2025), between 15:00 and 17:00, the first leaf was wounded on both sides (six scratches per half leaf using serrated forceps) and the entire wounded surface treated with 8 µL of regurgitant. The second leaf was wounded on one side and treated with 4 µL. On the day of measurements (2 May 2025), induction of the first row was performed between 07:30 and 08:30 and induction of the second row between 09:30 and 10:00. On this second day, the remaining half of leaf 2 was wounded and treated with 4 µL, and leaf 3 was wounded on both sides and treated with 8 µL. As a result, plants were sampled approximately 4–5 h after the most recent induction.

#### Volatile measurements

volatile emissions were measured individually for each plant on 2 May 2025 between 11:15 and 14:45 (n= 67 damaged, n=64 control). Within each treatment group, one plant located at the center of the group and one at the border were randomly selected for sampling. Sampling progressed systematically along the rows, starting with the left row and then moving to the right row. Meteorological variables were automatically recorded every 2 min using a small weather station (DNT Pro, GmbH & Co. KG, Germany). During the sampling time, solar radiation ranged from 343.4 to 708.2 W m⁻² under clear, sunny conditions, air temperature ranged from 22.5 to 27.5 °C, and relative humidity from 34 to 54%. Wind speeds were low, ranging from 0 to 2.6 m s⁻¹, corresponding to light wind conditions. However, wind direction was highly variable throughout the measurement period (2–368°, median 257°), preventing consistent downwind positioning as achieved in the simpler open-air experiment. Sampling was performed from one of the sides of the rows and kept constant for each row. Based on recorded wind direction and sampling side, the angular difference between approximate inlet orientation and airflow direction (0–180°, where 0° indicates downwind and 180° upwind sampling) had a median of 125° (range 30–180°), reflecting variable and often non-optimal sampling geometry relative to wind. Because inlet positioning was approximate and a 10 cm diameter funnel was used to broaden the sampling cone, inlet–wind alignment should be considered indicative rather than exact.

### Data analysis

Statistical analyses and modeling were performed using a combination of R (4.3.2; R Core Team, 2023) and Python (version 3.10.12). Classical statistical analyses and Random Forest modeling for laboratory assays were primarily conducted in R, whereas machine-learning analyses for open-air and field measurements were performed in Python using the scikit-learn library (version 1.5.0; Pedregosa et al., 2011).

#### Laboratory measurements

To assess the discriminant capacity of the three odor-sensing methods under controlled conditions, we applied the same multivariate workflow to all laboratory datasets. We performed non-metric multidimensional scaling (NMDS) and a permutational multivariate analysis of variance (PERMANOVA) (vegan; Oksanen et al., 2022) to assess treatment-level separation of overall volatile profiles. NMDS was computed on a Gower dissimilarity matrix and constrained to two dimensions. In parallel, univariate treatment effects were assessed for each quantified compound (GC-MS) or targeted ion (PTR-TOF) using heteroscedasticity-robust ANOVAs, implemented with the *Anova* function (car; Fox and Weisberg, 2019) in combination with heteroscedasticity-consistent covariance estimators (sandwich; Zeileis et al., 2019). Post-hoc comparisons were performed using *glht* (multcomp; Hothorn et al., 2016) with false discovery rate (FDR) correction. To test whether volatile profiles could predict treatment, we used Random Forest (RF) classification implemented in the *randomForest* package (Liaw and Wiener, 2002). Model performance was evaluated using the out-of-bag (OOB) procedure, an internal validation method in which a subset of samples is automatically withheld during model training. Prediction performance was summarized using classification accuracy and confusion matrices, which describe the proportion of correctly classified samples and the distribution of predicted versus observed treatments.

#### Open-air measurements with PTR-TOF

All analyses were performed separately for each sampling distance (1–2 cm and 4–5 cm). Data structure was first explored by principal component analysis (PCA; scikit-learn) applied to ion concentrations averaged over the 60-s sampling period and standardized to zero mean and unit variance. This analysis allowed to visualize and assess variation among sampling days, across temperature and humidity gradients and between treatments. We used the humidity and temperature values recorded with the small weather station as described above. Then, to identify ions detected at higher levels in damaged than control plants, we performed Wilcoxon rank-sum tests on the 60-s averaged values.

To assess whether caterpillar damage could be predicted at the individual plant level from real-time odor profiles and to evaluate the influence of sampling duration, we implemented a supervised machine-learning workflow. Each 60-s measurement was partitioned into 1, 5, 15, or 30 s windows, and concentrations were aggregated by averaging within each window. Two dataset constructions were used: Dataset A, in which all time windows from a given plant were retained (yielding *n* × 60/*window length* observations per plant), and Dataset B, in which only the first time window of each measurement was retained, resulting in one observation per plant, thereby avoiding repeated measurements from the same individual and allowing direct comparison among sampling durations without effects of unequal sample sizes. Datasets were split into training (70%) and testing (30%) sets. To prevent leakage in Dataset A (non-independence among multiple windows from the same plant), splitting and cross-validation were grouped by plant, ensuring that all measurements from one plant remained in the same fold/split. Features (ions) were always to zero mean and unit variance to put all features on a comparable scale.

We then trained and compared the following classifiers: logistic regression with L1 regularization (Lasso; saga solver), k-nearest neighbors, support vector machine, decision tree, and random forest. Performance was evaluated using five-fold cross-validation (grouped by plant for Dataset A). Models were compared by mean accuracy across folds, and the associated 95% confidence interval.

Because 1-s windows yielded high performance and L1-regularized logistic regression performed as well as or better than other algorithms, subsequent analyses focused on Dataset A (1 s windows). Moreover, L1 (Lasso) regularization limits model complexity by shrinking regression coefficients, with weakly informative features being reduced to zero. This allows feature selection, facilitates biological interpretation by identifying the most informative ions, and reduces overfitting, thereby improving model robustness and generalizability. The regularization strength (hyperparameter C) was tuned using grouped five-fold cross-validation over 10 values from 0.001 to 1. The selected values were C = 0.223 (1–2 cm) and C = 0.112 (4–5 cm). Final models were evaluated on the test set using confusion matrices and reporting accuracy. Model coefficients (weights) were used to assess feature importance, with larger absolute coefficient values indicating ions that contributed most strongly to the discrimination between damaged and control plants.

Finally, to test robustness to day-to-day variability, we additionally performed a leave-one-day-out evaluation: models were trained on two sampling days and tested on the third day, for all combinations and for both distances, using default C values.

#### Open-air measurements with MSS

A PCA was performed on sensor responses extracted from the 6th sampling phase to visualize variation among treatments, sampling days, and gradients of temperature and humidity. For these analyses, temperature and humidity were obtained from the internal sensor of the MSS module. Multiple feature extraction strategies from MSS response curves and alternative preprocessing approaches were evaluated, including the laboratory strategy (features extracted from one sampling phase) and previously described MSS workflows (Lang et al., 2016; Saeki et al., 2024). For each processed dataset, the same machine-learning algorithms as above were evaluated using k-fold cross-validation, and performance was compared to a dummy-classifier baseline (random prediction and majority-class prediction).

#### Field measurements with ultra-portable TOF

In contrast to previously analyzed PTR-TOF datasets, which relied on fitted peak areas quantified in absolute concentration units, the present analyses employed raw ion counts aggregated for each nominal m/z from 7 to 556, summing all counts within ±0.5 Da of each nominal mass. No targeted peak fitting was applied, allowing an exploratory analysis to identify spectral markers relevant to plant damage under field conditions.

For each 60-second measurement, the average aggregated intensity per m/z value was used. To evaluate the potential of the portable TOF measurements to discriminate between damaged and undamaged plants and to identify the most informative m/z features, we applied logistic regression with L1 (Lasso) regularization (saga solver). This method was chosen because it performed well in previous open-air analyses and performs embedded feature selection while handling high-dimensional data with potentially irrelevant or collinear features. Prior to modeling, the raw intensities were probabilistic quotient normalized (PQN) using the median spectrum as reference to correct for differences in overall signal intensity between samples. This approach corrects sample-specific dilution effects while preserving the relative contribution of individual features. Intensities were then log-transformed (log1p) to reduce skew and stabilize variance and standardized to zero mean and unit variance to put all features on a comparable scale.

Because the dataset was relatively small, hyperparameters were left at default values, and no separate test set was held out for validation. Instead, repeated stratified five-fold cross-validation (ten repeats) was used to obtain robust estimates of model performance and feature stability. Metrics were reported as mean values across repeats with 95% confidence intervals. Feature importance was assessed based on the mean coefficient values (weights). Univariate statistical tests (Wilcoxon rank-sum tests) were additionally performed to assess differences in ion intensities between treatments for the most informative features.

## Data and materials availability

Data and codes are available at https://github.com/ceco-lab/sensor.git

## Supplementary information

see Supplementary_Material_Data.pdf

## Acknowledgements

We thank our supportive colleagues in the FARCE group, including Christèle Borgeaud, Kathrin Altermatt, Kilian Alemano, Vladimir Redondo and Ilham Jouini at the University of Neuchâtel for helping with insect rearing, plant growing, laboratory maintenance and helping during the open-air and field measurements. We also thank Gabriele Macgregor from Tofwerk for helping during the initial tests of the portable TOF instrument. We also thank Ecorobotix for providing and preparing the maize plot for the field experiment, in particular Elise Mettraux and Pauline Anne. This work was supported by the University of Neuchâtel and advanced grant no. 788949 of the European Research Council awarded to T.C.J.T and in part by Horizon Europe project PurPest, grant no. 101060634. This study was also partially supported by the Public/Private R&D Investment Strategic Expansion Program (PRISM), Cabinet Office, Japan.

## Author contributions

T.C.J.T., C.C.M.A., M.M., G.R., A.K., conceived and designed the project. C.C. M.A., M.M., G.R., A.K. performed laboratory measurements. C.C.M.A., M.M., A.K. G.R. performed open-air measurements. M.M., C.C.M.A., F.L., P.B., performed field measurements. T.A., K.M., G.Y., F.L., L.C. designed sensor equipment and/or provided technical advice. M.M., E.D., C.C.M.A., G.R., A.K., S.R., T.D, Y.L. conducted and advised on data analyses. M.M., C.C.M.A, G.R., T.C.J.T. wrote the manuscript, with contributions from all authors.

## Supplementary Materials for

**Fig. S1.**
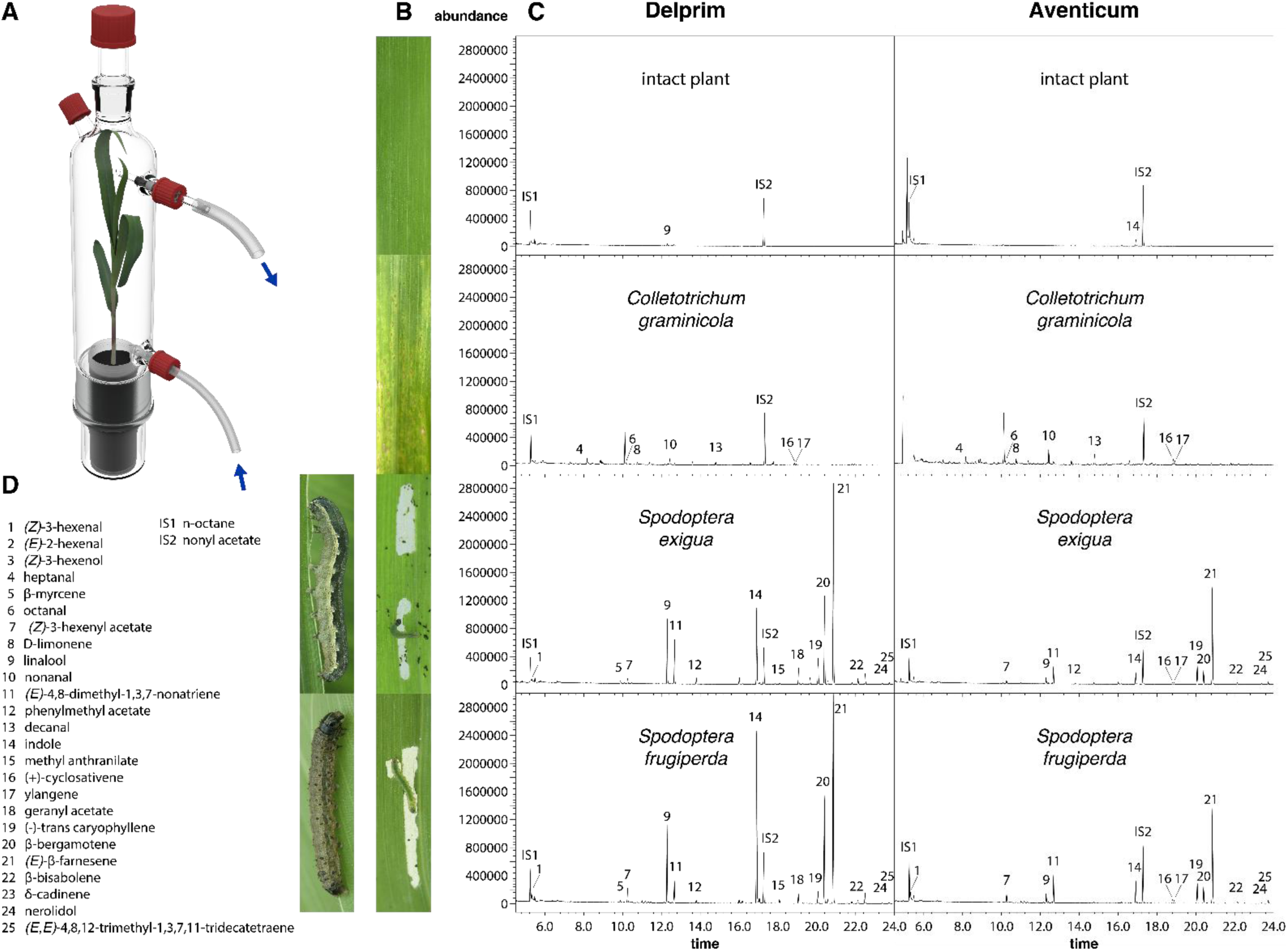
Volatile profiles of maize plants infested by caterpillars and a fungal pathogen. (A) Dynamic headspace sampling setup used to collect volatile emissions from maize plants enclosed in glass vessels. (B) Pictures of the different treatments, showing an undamaged leaf and leaves damaged by two herbivore species or infected by a fungal pathogen. (C) Examples of gas chromatography–mass spectrometry (GC–MS) chromatograms (total ion current) obtained from the different treatments. (D) List of the main volatile compounds detected across treatments.

**Fig. S2.**
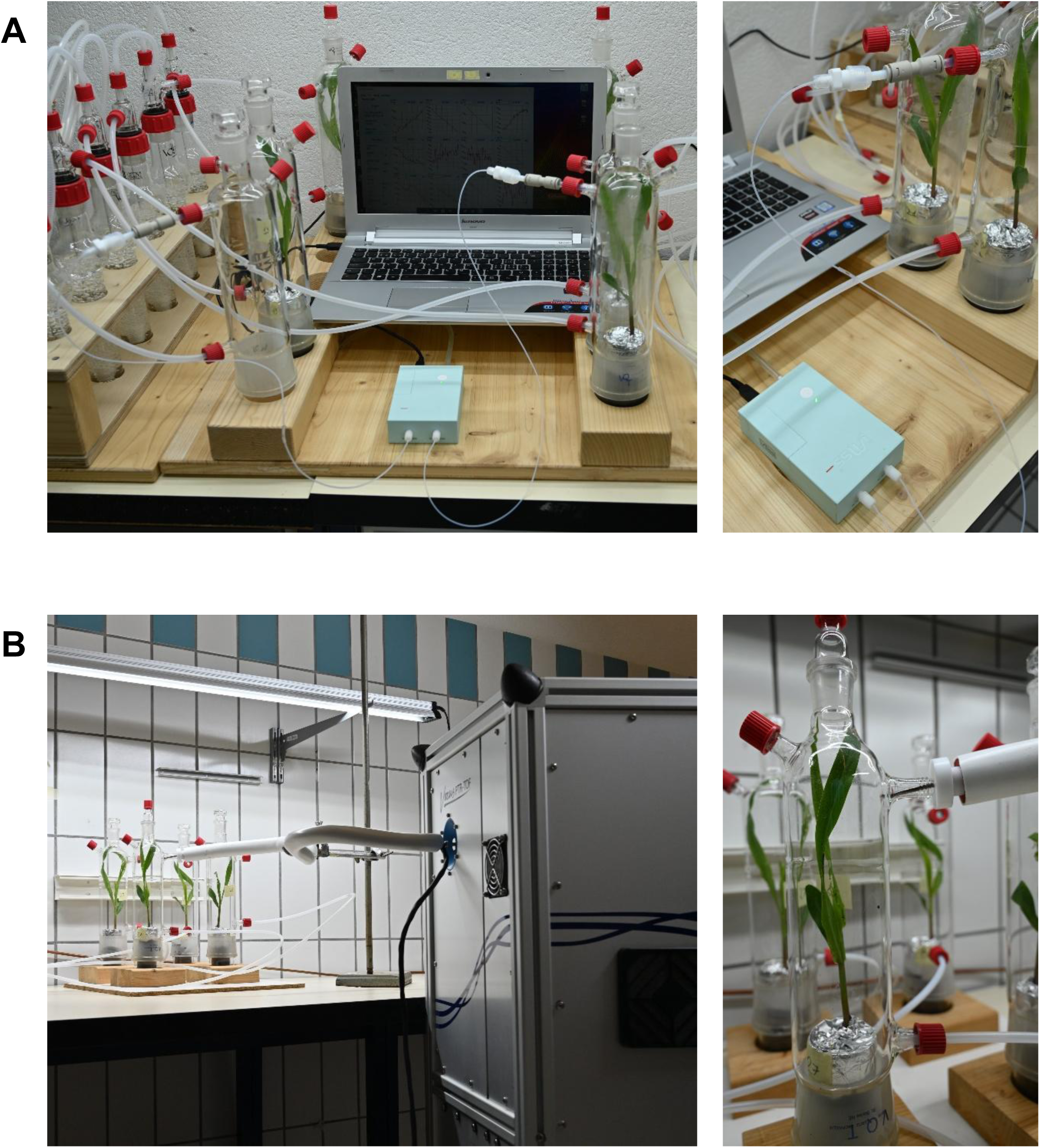
Dynamic headspace sampling setups used for laboratory measurements of maize volatile emissions. (A) Sampling configuration coupled to the membrane-type surface stress sensor (MSS) module. (B) Sampling configuration coupled to a proton-transfer-reaction time-of-flight (PTR-TOF) mass spectrometer (Vocus S). *Photographs were taken by the authors*.

**Fig. S3.**
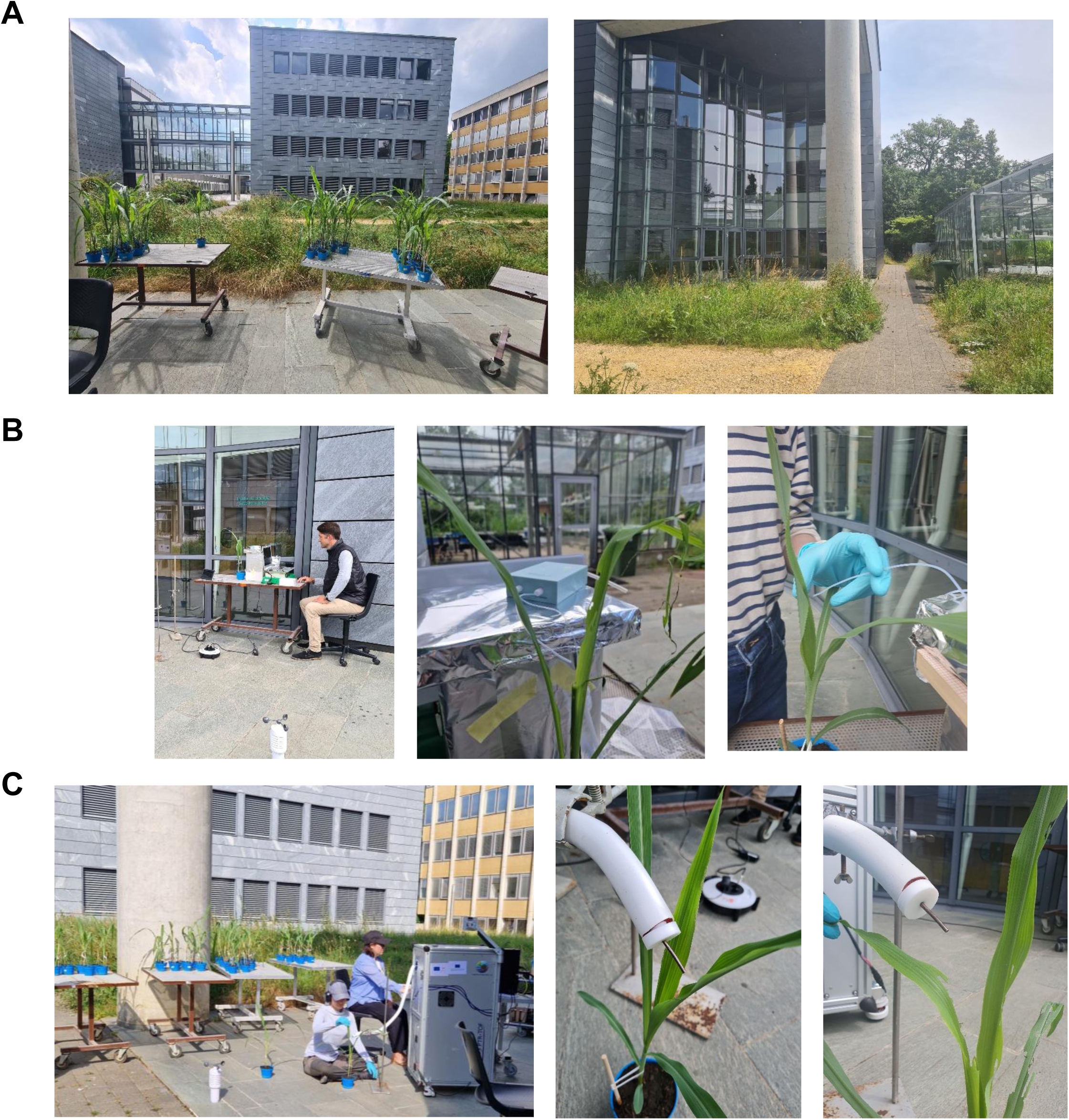
Outdoor open-air sampling setup on campus for MSS module and PTR-TOF (Vocus S) measurements of maize volatile emissions. Overview of the outdoor experimental setup used for open-air measurements conducted on campus. **(A)** Surrounding environment at the measurement site. **(B)** Measurement configuration using the membrane-type surface stress sensors (MSS) module. **(C)** Measurement configuration using the PTR-TOF (Vocus S) instrument. *Photographs depict the authors, who provided consent for publication*.

**Fig. S4.**
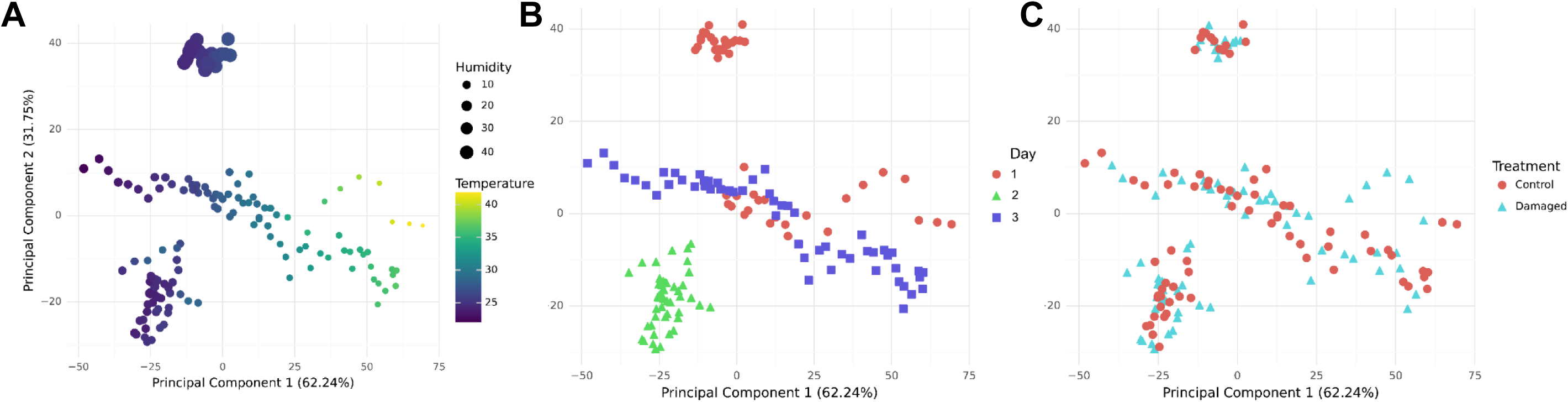
PCA of MSS module response recorded during outdoor open-air sampling of volatiles from undamaged and *Spodoptera exigua*–damaged maize plants over three sampling days. The first two principal components are shown. Sensor features were extracted from the sampling phase of the sixth measurement cycle (110–120 s), subsampling every 10th data point, and standardized to zero mean and unit variance prior to analysis. **(A)** Scores colored by ambient temperature, with point size indicating relative humidity. **(B)** Scores colored and shaped according to sampling day. **(C)** Scores colored and shaped according to plant treatment (undamaged vs. damaged).

**Fig. S5.**
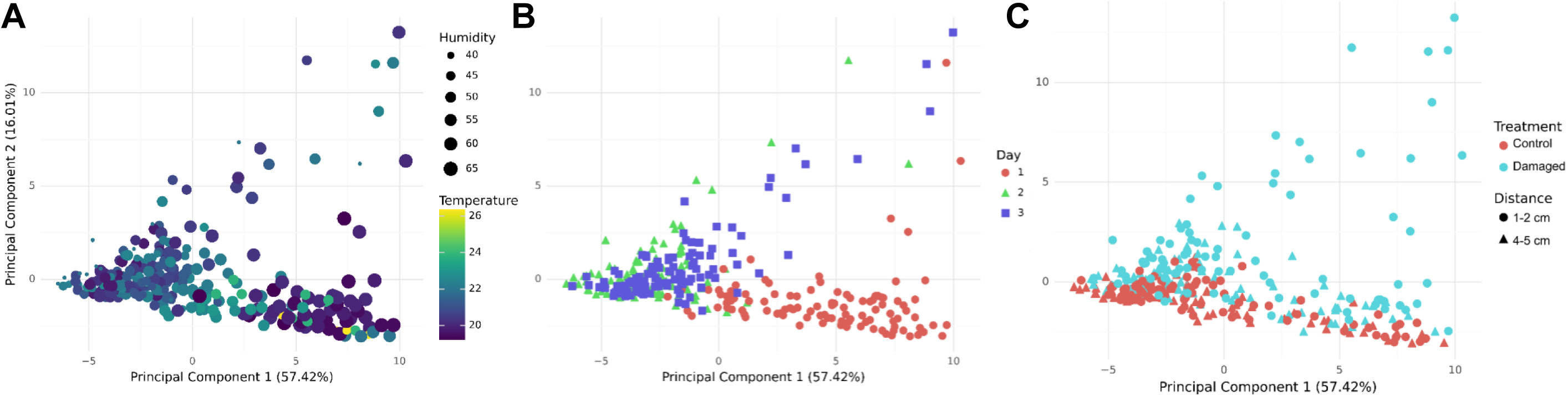
PCA of PTR-TOF (Vocus S) measurements recorded during outdoor open-air sampling of volatiles from undamaged and *Spodoptera exigua*–damaged maize plants over three sampling days. The first two principal components are shown. Analyses were performed on concentrations averaged over the 60-s sampling period and standardized to zero mean and unit variance prior to analysis. **(A)** Scores colored by ambient temperature, with point size indicating relative humidity. **(B)** Scores colored and shaped according to sampling day. **(C)** Scores colored according to plant treatment (undamaged vs. damaged) and shaped according to sampling distance (1–2 cm vs. 4–5 cm).

**Fig. S6.**
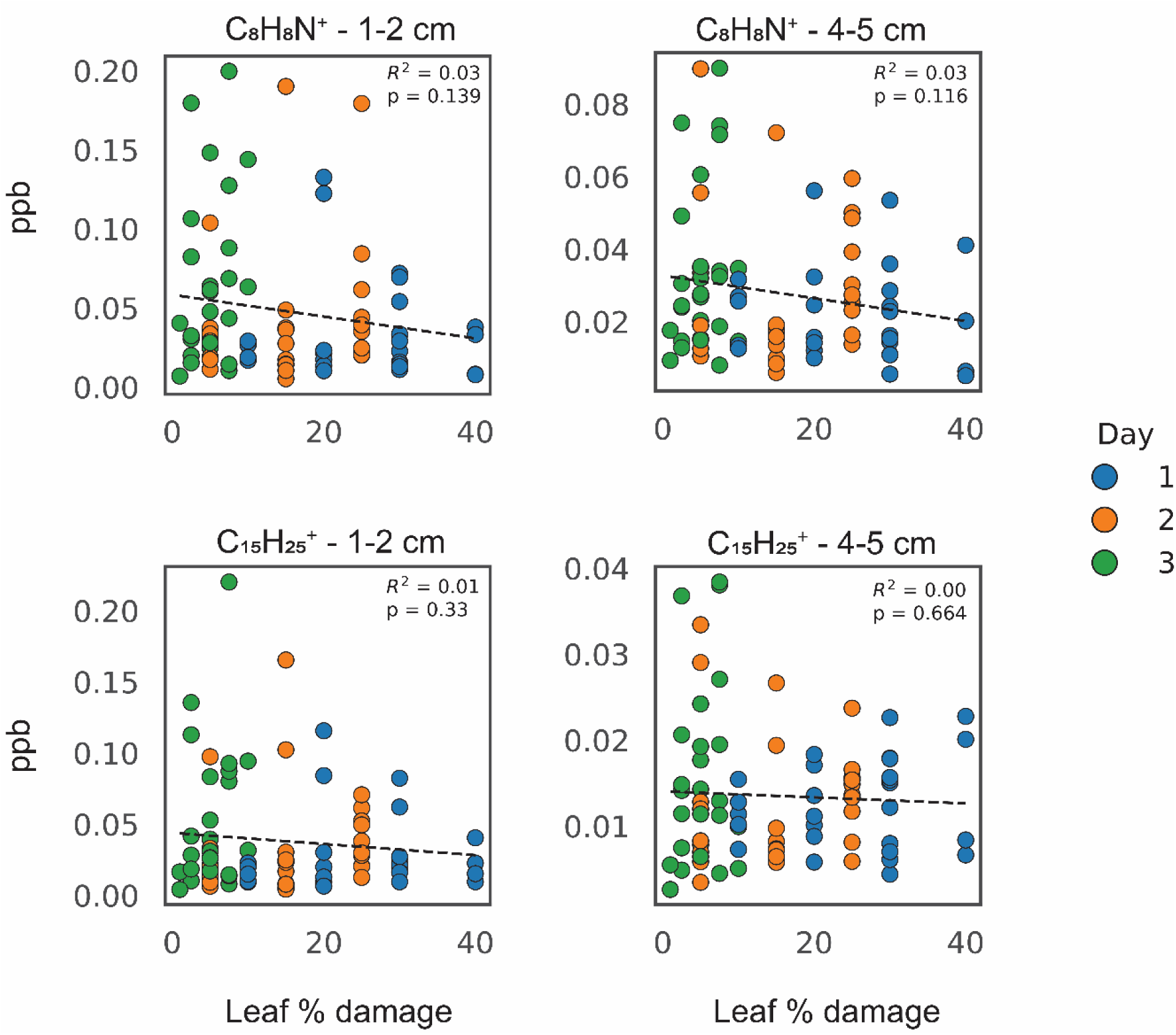
Relationship between visually assessed leaf percentage of damage and volatile concentrations measured by PTR-TOF (Vocus S) during outdoor open-air sampling. Linear regression analyses of concentrations of indole (C₈H₈N⁺) and sesquiterpenes (C₁₅H₂₅⁺) measured by PTR-TOF in *Spodoptera exigua*–damaged maize plants. Data are shown separately for measurements taken closer to the leaves (1–2 cm) and farther from the leaves (4–5 cm). Each point represents concentrations averaged over a 60-s measurement and is colored according to the day of sampling. Dashed lines indicate linear regression fits; coefficients of determination (R²) and associated p-values from the linear regression are shown on each panel.

**Fig. S7.**
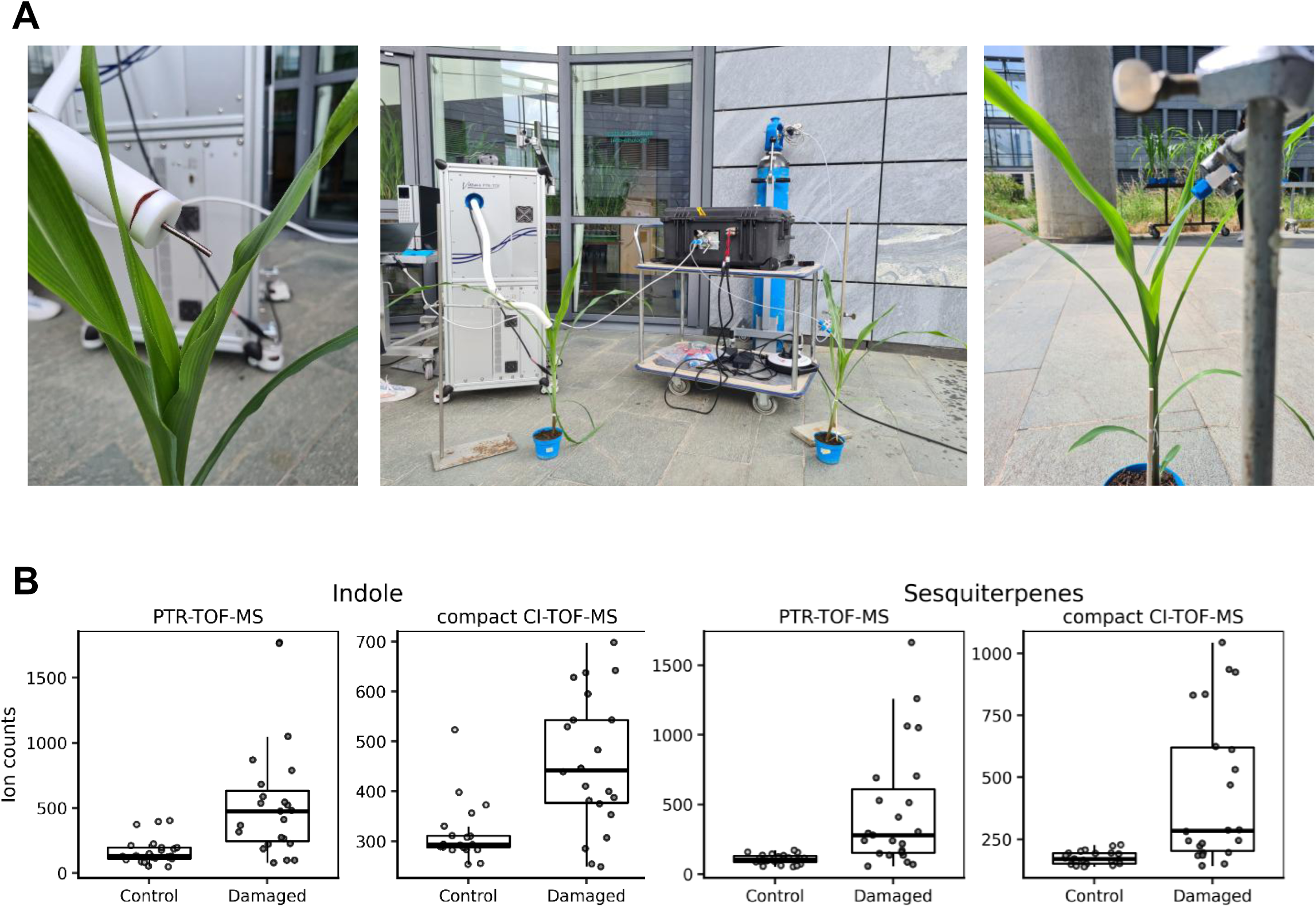
Evaluation in comparison to PTR-TOF (Vocus S) of the ultra-portable TOF (Vocus C) instrument for detection of herbivory-associated volatiles in maize in open-air conditions. **(A)** Photographs illustrating the experimental setup used for comparison of real-time mass spectrometry instruments prior to field assays. The central image shows the PTR-TOF instrument on the left and the portable TOF on the right, operated using benzene-adduct cation ionization. The left close-up image shows the PTR-TOF sampling inlet positioned near maize leaves subjected to simulated herbivory, while the right close-up image shows the portable TOF sampling inlet. The same plants were measured successively with both instruments. (B) Comparison of herbivory-associated volatile signals measured by PTR-TOF and portable TOF during open-air measurements on greenhouse-grown maize plants exposed to simulated caterpillar damage. Faceted boxplots show ion counts for indole and sesquiterpenes in control and damaged plants. Indole was detected as C₈H₈N⁺ by PTR-TOF and at m/z 117 by portable TOF, whereas sesquiterpenes were detected as C₁₅H₂₅⁺ by PTR-TOF and at m/z 204 by compact portable TOF. Statistical comparisons between control and damaged plants were performed using Wilcoxon rank-sum tests (indole, PTR-TOF: W = 82, p < 0.001; indole, portable TOF: W = 81, p < 0.001; sesquiterpenes, PTR-TOF: W = 69, p < 0.001; sesquiterpenes, portable TOF: W = 69, p < 0.001). Sample sizes were n = 45 damaged plants and n = 43 control plants. *Photographs were taken by the authors*.

**Table S1.** Emission rates of volatile compounds emitted by maize plants (var. *Delprim*) measured by GC–MS under laboratory conditions. Emission rates are shown as mean ± s.e. (ng h⁻¹) for undamaged control plants and plants damaged by six *Spodoptera exigua larvae*, damaged by four *Spodoptera frugiperda* larvae, or infected by *Colletotrichum graminicola* (n = 7–8 plants per treatment). Volatiles were collected 24 h after the onset of herbivory and 7 days after fungal inoculation and analyzed by gas chromatography–mass spectrometry (GC–MS). F-statistics and p-values from robust analyses of variance (ANOVA) are reported. Different letters indicate significant pairwise differences after false discovery rate (FDR) correction using a heteroscedasticity-consistent covariance estimator.

| Var. *Delprim* emission rate (ng h<sup>-1</sup>) |  |  |  |  |  |  |
| --- | --- | --- | --- | --- | --- | --- |
| Compounds | Control | *C. graminicola* | *S. exigua* | *S. frugiperda* | F-stat. | P-value |
| (Z)-3-Hexenal | 0 ± 0 a | 0 ± 0 a | 246.56 ± 118.32 ab | 546.49 ± 90.75 b | 11.58 | < 0.001 |
| (E)-2-Hexenal | 0 ± 0 | 69.05 ± 25.29 | 0 ± 0 | 0 ± 0 | 2.17 | 0.116 |
| (Z)-3-Hexenol | 0 ± 0 a | 58.28 ± 8.47 b | 0 ± 0 a | 0 ± 0 a | 13.59 | < 0.001 |
| Heptanal | 0 ± 0 a | 70.48 ± 8.78 b | 0 ± 0 a | 0 ± 0 a | 18.52 | < 0.001 |
| β-Myrcene | 0 ± 0 a | 35.18 ± 7.47 b | 58.44 ± 10.86 b | 55.01 ± 6.76 b | 32.93 | < 0.001 |
| Octanal | 0 ± 0 | 736.85 ± 352.76 | 0 ± 0 | 0 ± 0 | 1.3 | 0.296 |
| (Z)-3-Hexenyl acetate | 0 ± 0 a | 148.94 ± 12.60 b | 247.21 ± 42.46 b | 235.97 ± 62.24 b | 54.19 | < 0.001 |
| D-Limonene | 0 ± 0 a | 166.64 ± 50.39 b | 0 ± 0 a | 0 ± 0 a | 3.2 | 0.04 |
| Linalool | 198.57 ± 63.12 a | 103.93 ± 38.86 a | 1670.52 ± 232.97 b | 1467.22 ± 227.52 b | 21.97 | < 0.001 |
| Nonanal | 0 ± 0 a | 269.63 ± 36.71 b | 0 ± 0 a | 0 ± 0 a | 15.69 | < 0.001 |
| DMNT | 0 ± 0 a | 325.71 ± 86.13 b | 714.00 ± 209.06 b | 397.02 ± 86.14 b | 13.54 | < 0.001 |
| Phenylmethyl acetate | 0 ± 0 a | 0 ± 0 a | 86.787 ± 18.81 b | 36.58 ± 7.19 b | 13.2 | < 0.001 |
| Decanal | 0 ± 0 a | 157.47 ± 14.91 b | 0 ± 0 a | 0 ± 0 a | 32.31 | < 0.001 |
| Indole | 0 ± 0 a | 14.83 ± 1.56 b | 2256.78 ± 178.56 c | 1205.28 ± 318.42 d | 72.83 | < 0.001 |
| Methyl anthranilate | 0 ± 0 a | 0 ± 0 a | 36.33 ± 4.59 b | 23.20 ± 6.80 b | 20.6 | < 0.001 |
| (+)-Cycloisosativene | 0 ± 0 | 10.86 ± 5.94 | 0 ± 0 | 0 ± 0 | 0.91 | 0.452 |
| Ylangene | 0 ± 0 a | 16.74 ± 4.97 b | 0 ± 0 a | 0 ± 0 a | 3.12 | 0.043 |
| Geranyl acetate | 0 ± 0 a | 0 ± 0 a | 275.84 ± 54.72 b | 165.52 ± 32.41 b | 14.64 | < 0.001 |
| β-Caryophyllene | 0 ± 0 a | 3.79 ± 1.60 a | 371.71 ± 57.00 b | 100.55 ± 24.94 c | 17.93 | < 0.001 |
| β-Bergamotene | 0 ± 0 a | 0 ± 0 a | 1261.48 ± 202.60 b | 816.83 ± 250.64 b | 14.1 | < 0.001 |
| (E)-β-Farnesene | 0 ± 0 a | 16.86 ± 1.19 b | 2908.32 ± 446.72 c | 1721.24 ± 544.20 c | 68.57 | < 0.001 |
| β-Bisabolene | 0 ± 0 a | 0 ± 0 a | 67.48 ± 13.69 b | 25.08 ± 8.77 ab | 9.08 | < 0.001 |
| δ-Cadinene | 0 ± 0 | 1.46 ± 0.96 | 0 ± 0 | 0 ± 0 | 0.29 | 0.833 |
| Nerolidol | 0 ± 0 a | 0 ± 0 a | 36.37 ± 10.03 b | 8.15 ± 5.40 ab | 4.22 | 0.015 |
| TMTT | 0 | 0 | 0 | 0 | - | - |
| Total | 198.57 ± 63.12 a | 2206.72 ± 392.26 b | 10237.84 ± 1305.85 c | 6804.16 ± 1606.29 bc | 28.85 | < 0.001 |

**Table S2.** Emission rates of volatile compounds emitted by maize plants (var. *Aventicum*) measured by GC–MS under laboratory conditions. Emission rates are shown as mean ± s.e. (ng h⁻¹) for undamaged control plants and plants damaged by six *Spodoptera exigua* larvae, damaged by four *Spodoptera frugiperda* larvae, or infected by *Colletotrichum graminicola* (n = 7–9 plants per treatment). Volatiles were collected 24 h after the onset of herbivory and 7 days after fungal inoculation and analyzed by gas chromatography–mass spectrometry (GC–MS). F-statistics and p-values from robust analyses of variance (ANOVA) are reported. Different letters indicate significant pairwise differences after false discovery rate (FDR) correction using a heteroscedasticity-consistent covariance estimator.

| Var. *Aventicum* emission rate (ng h<sup>-1</sup>) |  |  |  |  |  |  |
| --- | --- | --- | --- | --- | --- | --- |
| Compounds | Control | *C. graminicola* | *S. exigua* | *S. frugiperda* | F-stat. | P-value |
| (Z)-3-Hexenal | 0 | 0 | 0 | 0 | - | - |
| (E)-2-Hexenal | 0 ± 0 | 482.39 ± 221.03 | 0 ± 0 | 0 ± 0 | 1.43 | 0.254 |
| (Z)-3-Hexenol | 0 ± 0 | 576.34 ± 251.41 | 0 ± 0 | 0 ± 0 | 1.58 | 0.216 |
| Heptanal | 0 ± 0 a | 180.29 ± 24.02 b | 0 ± 0 a | 0 ± 0 a | 16.62 | < 0.001 |
| β-Myrcene | 0 ± 0 a | 138.39 ± 39.09 b | 0 ± 0 a | 0 ± 0 a | 3.71 | 0.023 |
| Octanal | 0 ± 0 | 1518.95 ± 544.86 | 0 ± 0 | 0 ± 0 | 2.33 | 0.096 |
| (Z)-3-Hexenyl acetate | 0 ± 0 a | 45.33 ± 21.89 a | 168.46 ± 31.17 b | 170.59 ± 33.69 b | 16.99 | < 0.001 |
| D-Limonene | 0 ± 0 a | 208.95 ± 39.56 b | 0 ± 0 a | 0 ± 0 a | 8.25 | < 0.001 |
| Linalool | 0 ± 0 a | 117.55 ± 31.46 b | 612.16 ± 116.32 c | 410.27 ± 125.80 bc | 15.2 | < 0.001 |
| Nonanal | 0 ± 0 a | 715.62 ± 112.49 b | 0 ± 0 a | 0 ± 0 a | 12 | < 0.001 |
| DMNT | 0 ± 0 a | 897.32 ± 239.31 b | 1482.01 ± 258.47 b | 840.04 ± 203.83 b | 18.59 | < 0.001 |
| Phenylmethyl acetate | 0 ± 0 a | 0 ± 0 a | 85.34 ± 33.24 ab | 16.60 ± 4.30 b | 5.89 | < 0.01 |
| Decanal | 0 ± 0 a | 472.14 ± 109.54 b | 0 ± 0 a | 0 ± 0 a | 5.52 | < 0.01 |
| Indole | 126.85 ± 34.60 a | 51.54 ± 9.74 a | 742.31 ± 123.42 b | 402.03 ± 128.06 ab | 12.24 | < 0.001 |
| Methyl anthranilate | 0 ± 0 | 0 ± 0 | 13.61 ± 6.02 | 0 ± 0 | 1.4 | 0.264 |
| (+)-Cycloisosativene | 0 ± 0 a | 61.44 ± 14.48 b | 35.00 ± 6.22 b | 34.89 ± 7.49 b | 20.22 | < 0.001 |
| Ylangene | 0 ± 0 a | 34.62 ± 11.58 b | 30.09 ± 5.05 b | 26.81 ± 5.88 b | 18.24 | < 0.001 |
| Geranyl acetate | 0 ± 0 | 0 ± 0 | 16.54 ± 5.92 | 0 ± 0 | 2.15 | 0.116 |
| β-Caryophyllene | 0 ± 0 a | 7.81 ± 3.81 a | 636.98 ± 114.96 b | 453.43 ± 181.77 ab | 11.8 | < 0.001 |
| β-Bergamotene | 0 ± 0 a | 0 ± 0 a | 505.09 ± 72.34 b | 421.49 ± 164.07 ab | 16.08 | < 0.001 |
| (E)-β-Farnesene | 0 ± 0 a | 60.60 ± 7.63 b | 3170.74 ± 396.04 c | 2526.72 ± 991.77 abc | 38.92 | < 0.001 |
| β-Bisabolene | 0 ± 0 a | 0 ± 0 a | 40.35 ± 6.63 b | 25.444 ± 10.95 ab | 12.01 | < 0.001 |
| δ-Cadinene | 0 ± 0 | 3.77 ± 1.40 | 0 ± 0 | 0 ± 0 | 1.61 | 0.208 |
| Nerolidol | 0 ± 0 a | 0 ± 0 a | 88.14 ± 18.64 b | 21.75 ± 11.81 a | 7.38 | < 0.01 |
| TMTT | 0 ± 0 a | 0 ± 0 a | 124.10 ± 20.86 b | 79.76 ± 26.30 b | 12.84 | < 0.001 |
| Total | 126.85 ± 34.60 a | 5573.07 ± 641.97 b | 7750.91 ± 1062.67 b | 5429.83 ± 1750.78 b | 38.79 | < 0.001 |

**Table S3.** Coating materials used in the membrane-type surface stress sensors (MSS) module. List of adsorbent polymers and nanoparticles coated on the 12 individual sensing channels of the piezoresistive membrane-type surface stress sensors (MSS) module.

|  | Channels | Coating |
| --- | --- | --- |
| **Array 1** | CH1 | Poly (2,6-Diphenyl-p-phenylene Oxide) (Tenax) |
|  | CH2 | Octadecyl modified silica–titania hybrid nanoparticles (C18-STNPs) |
|  | CH3 | Phenyl modified silica–titania hybrid nanoparticles (Ph-STNPs) |
|  | CH4 | Poly (4-methyl styrene) (P4MS) |
| **Array 2** | CH5 | Polycaprolactone (PCL) |
|  | CH6 | Polystyrene (PS) |
|  | CH7 | Polyvinyl fluoride (PVF) |
|  | CH8 | Poly (methyl methacrylate) (PMMA) |
| **Array 3** | CH9 | Poly (methyl vinyl ether-alt-maleic anhydride) (PMVE) |
|  | CH10 | Phenyl modified silica–titania hybrid nanoparticles (Ph-STNPs) |
|  | CH11 | Poly (ethyleneimine) (PEI) |
|  | CH12 | Propyleneglycol Adipate (PGA) |

**Table S4.**
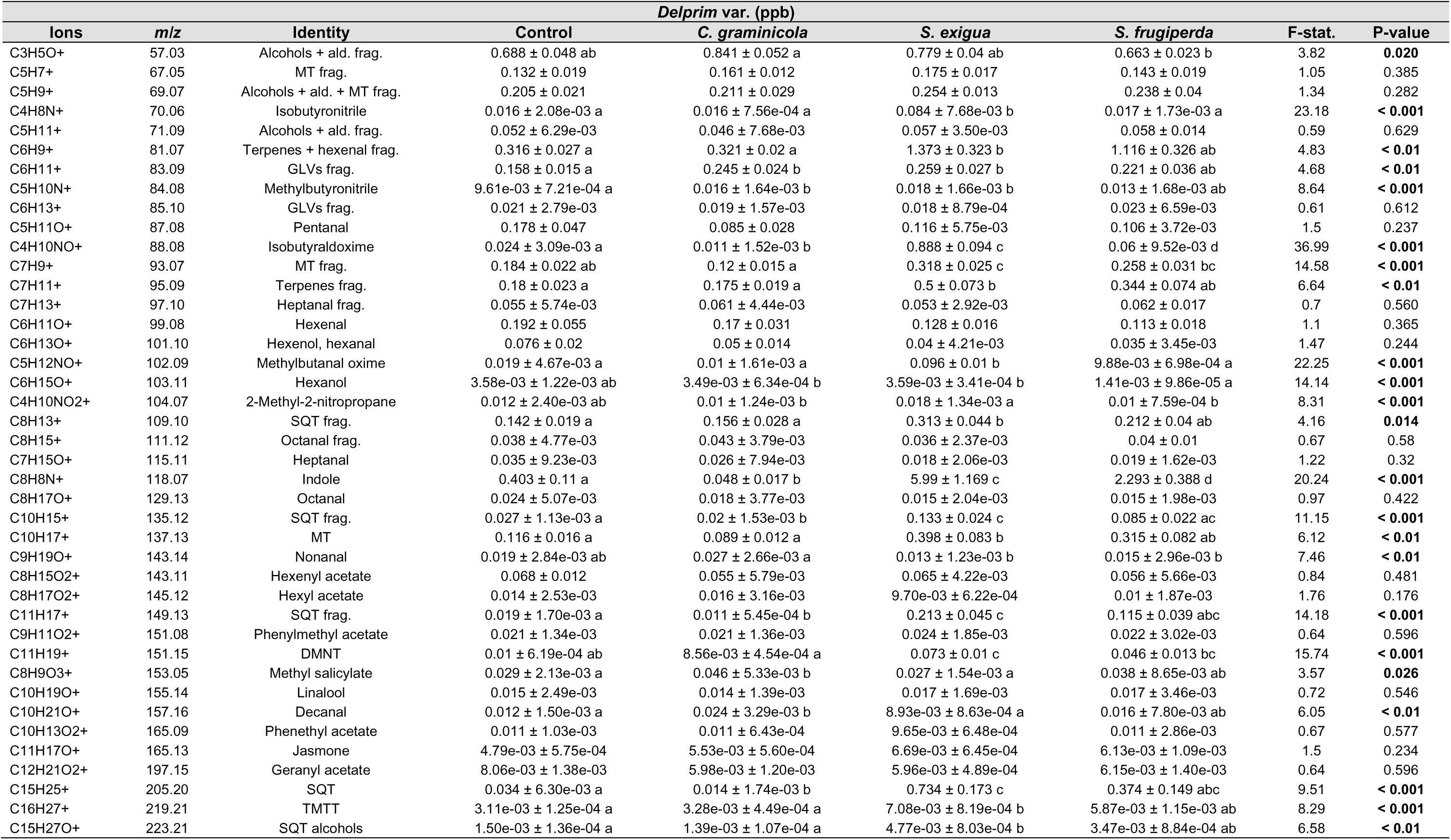
Concentrations of volatile compounds (protonated ions) detected by PTR-TOF (Vocus S) from laboratory headspace samples of maize plants (var. *Delprim*). Concentrations are shown as mean ± s.e. (ppb) for undamaged plants and plants infected by *Colletotrichum graminicola* or damaged by *Spodoptera exigua* or *S. frugiperda* larvae (n = 5–8 plants per treatment). Volatile emissions were measured 24 h after the onset of herbivory and 7 days after fungal inoculation using PTR-TOF. Values correspond to concentrations averaged over 60-s measurements (1 spectrum s⁻¹). Reported ions represent protonated volatile compounds (parent ions) or fragment ions. F-statistics and p-values from robust analyses of variance (ANOVA) are reported, and different letters indicate significant pairwise differences after false discovery rate (FDR) correction using a heteroscedasticity-consistent covariance estimator. MT, monoterpenes; SQT, sesquiterpenes; ald., aldehyde; frag., fragment.

**Table S5.** Concentrations of volatile compounds (protonated ions) detected by PTR-TOF (Vocus S) from laboratory headspace samples of maize plants (var. *Aventicum*). Concentrations are shown as mean ± s.e. (ppb) for undamaged plants and plants infected by *Colletotrichum graminicola* or damaged by *Spodoptera exigua* or *S. frugiperda* larvae (n = 5–8 plants per treatment). Volatile emissions were measured 24 h after the onset of herbivory and 7 days after fungal inoculation using PTR-TOF. Values correspond to concentrations averaged over 60-s measurements (1 spectrum s⁻¹). Reported ions represent protonated volatile compounds (parent ions) or fragment ions. F-statistics and p-values from robust analyses of variance (ANOVA) are reported, and different letters indicate significant pairwise differences after false discovery rate (FDR) correction using a heteroscedasticity-consistent covariance estimator. MT, monoterpenes; SQT, sesquiterpenes; frag., fragment

| Ions | m/z | Attribution | Aventicum var. (ppb) |  |  |  | F-stat. | P-value |
| --- | --- | --- | --- | --- | --- | --- | --- | --- |
|  |  |  | Control | *C. graminicola* | *S. exigua* | *S. frugiperda* |  |  |
| C3H5O+ | 57.03 | Alcohols + ald. frag. | 1.047 ± 0.123 | 0.943 ± 0.164 | 1.276 ± 0.148 | 1.172 ± 0.247 | 0.73 | 0.540 |
| C5H7+ | 67.05 | MT frag. | 0.139 ± 0.015 a | 0.162 ± 0.018 a | 0.265 ± 0.024 b | 0.201 ± 0.028 ab | 6.19 | < 0.01 |
| C5H9+ | 69.07 | Alcohols + ald. + MT frag. | 0.25 ± 0.037 | 0.232 ± 0.066 | 0.41 ± 0.05 | 0.316 ± 0.067 | 2.29 | 0.100 |
| C4H8N+ | 70.06 | Isobutyronitrile | 0.023 ± 4.72e-03 ab | 0.016 ± 2.66e-03 a | 0.182 ± 0.058 b | 0.04 ± 0.01 ab | 4.02 | 0.016 |
| C5H11+ | 71.09 | Alcohols + ald. frag. | 0.19 ± 0.138 | 0.055 ± 0.015 | 0.184 ± 0.112 | 0.366 ± 0.312 | 0.94 | 0.435 |
| C6H9+ | 81.07 | Terpenes + hexenal frag. | 0.34 ± 0.117 ab | 0.25 ± 0.047 a | 1.816 ± 0.277 c | 1.836 ± 0.529 bc | 11.47 | < 0.001 |
| C6H11+ | 83.09 | GLVs frag. | 0.276 ± 0.061 | 0.302 ± 0.111 | 0.361 ± 0.073 | 0.36 ± 0.095 | 0.31 | 0.819 |
| C5H10N+ | 84.08 | Methylbutyronitrile | 0.019 ± 4.81e-03 | 0.02 ± 7.64e-03 | 0.035 ± 7.45e-03 | 0.025 ± 7.70e-03 | 0.98 | 0.417 |
| C6H13+ | 85.10 | GLVs frag. | 0.045 ± 0.026 | 0.022 ± 3.94e-03 | 0.054 ± 0.035 | 0.073 ± 0.055 | 0.73 | 0.543 |
| C5H11O+ | 87.08 | Pentanal | 0.21 ± 0.089 | 0.115 ± 0.042 | 0.237 ± 0.117 | 0.287 ± 0.168 | 0.68 | 0.574 |
| C4H10NO+ | 88.08 | Isobutyraldoxime | 0.045 ± 0.015 ab | 0.012 ± 1.85e-03 a | 2.386 ± 0.91 ab | 0.255 ± 0.076 b | 6.43 | < 0.01 |
| C7H9+ | 93.07 | MT frag. | 0.232 ± 0.059 | 0.151 ± 0.066 | 0.365 ± 0.061 | 0.305 ± 0.091 | 1.72 | 0.184 |
| C7H11+ | 95.09 | Terpenes frag. | 0.134 ± 0.015 a | 0.162 ± 0.023 a | 0.81 ± 0.194 b | 0.45 ± 0.121 ab | 5.58 | < 0.01 |
| C7H13+ | 97.10 | Heptanal frag. | 0.126 ± 0.062 | 0.065 ± 0.017 | 0.155 ± 0.088 | 0.174 ± 0.116 | 0.74 | 0.534 |
| C6H11O+ | 99.08 | Hexenal | 0.102 ± 0.016 | 0.153 ± 0.013 | 0.119 ± 0.019 | 0.198 ± 0.082 | 2.08 | 0.124 |
| C6H13O+ | 101.10 | Hexenol, hexanal | 0.098 ± 0.05 | 0.051 ± 0.011 | 0.139 ± 0.099 | 0.139 ± 0.095 | 0.7 | 0.562 |
| C5H12NO+ | 102.09 | Methylbutanal oxime | 0.02 ± 6.32e-03 a | 9.87e-03 ± 1.64e-03 a | 0.393 ± 0.175 a | 0.061 ± 0.019 a | 4.11 | 0.015 |
| C6H15O+ | 103.11 | Hexanol | 0.014 ± 0.011 | 4.22e-03 ± 9.93e-04 | 0.025 ± 0.016 | 0.024 ± 0.021 | 0.97 | 0.418 |
| C4H10NO2+ | 104.07 | 2-Methyl-2-nitropropane | 0.013 ± 2.98e-03 | 0.01 ± 1.43e-03 | 0.028 ± 6.23e-03 | 0.015 ± 4.77e-03 | 2.54 | 0.076 |
| C8H13+ | 109.10 | SQT frag. | 0.17 ± 0.031 a | 0.133 ± 0.038 a | 0.481 ± 0.094 b | 0.33 ± 0.08 ab | 4.42 | 0.011 |
| C8H15+ | 111.12 | Octanal frag. | 0.115 ± 0.077 | 0.04 ± 7.69e-03 | 0.156 ± 0.113 | 0.166 ± 0.126 | 0.86 | 0.473 |
| C7H15O+ | 115.11 | Heptanal | 0.048 ± 0.022 | 0.028 ± 8.27e-03 | 0.063 ± 0.041 | 0.063 ± 0.041 | 0.55 | 0.650 |
| C8H8N+ | 118.07 | Indole | 0.54 ± 0.106 a | 0.054 ± 0.016 b | 2.551 ± 0.498 c | 1.133 ± 0.233 ac | 19.64 | < 0.001 |
| C8H17O+ | 129.13 | Octanal | 0.016 ± 1.80e-03 | 0.018 ± 3.27e-03 | 0.015 ± 3.13e-03 | 0.015 ± 2.82e-03 | 0.17 | 0.913 |
| C10H15+ | 135.12 | SQT frag. | 0.045 ± 7.69e-03 a | 0.02 ± 5.20e-03 a | 0.234 ± 0.028 b | 0.145 ± 0.034 b | 20.24 | < 0.001 |
| C10H17+ | 137.13 | MT | 0.042 ± 7.80e-03 a | 0.045 ± 0.019 a | 0.281 ± 0.056 b | 0.155 ± 0.047 ab | 6.83 | < 0.01 |
| C9H19O+ | 143.14 | Nonanal | 0.02 ± 4.43e-03 | 0.026 ± 7.93e-03 | 0.014 ± 2.50e-03 | 0.012 ± 2.08e-03 | 1.31 | 0.290 |
| C8H15O2+ | 143.11 | Hexenyl acetate | 0.044 ± 6.42e-03 | 0.046 ± 7.82e-03 | 0.051 ± 8.88e-03 | 0.05 ± 9.10e-03 | 0.16 | 0.921 |
| C8H17O2+ | 145.12 | Hexyl acetate | 0.02 ± 8.17e-03 | 0.015 ± 2.61e-03 | 0.024 ± 0.014 | 0.023 ± 0.013 | 0.28 | 0.842 |
| C11H17+ | 149.13 | SQT frag. | 0.028 ± 4.11e-03 a | 0.014 ± 1.96e-03 b | 0.331 ± 0.056 c | 0.21 ± 0.061 c | 14.89 | < 0.001 |
| C9H11O2+ | 151.08 | Phenylmethyl acetate | 0.023 ± 5.26e-03 | 0.026 ± 2.71e-03 | 0.028 ± 6.56e-03 | 0.027 ± 7.86e-03 | 0.17 | 0.916 |
| C11H19+ | 151.15 | DMNT | 0.012 ± 1.00e-03 a | 0.013 ± 3.41e-03 a | 0.248 ± 0.063 b | 0.1 ± 0.028 b | 7.11 | < 0.01 |
| C8H9O3+ | 153.05 | Methyl salicylate | 0.041 ± 5.62e-03 | 0.063 ± 0.014 | 0.037 ± 9.14e-03 | 0.04 ± 6.81e-03 | 0.7 | 0.560 |
| C10H19O+ | 155.14 | Linalool | 0.011 ± 1.20e-03 | 0.011 ± 1.71e-03 | 0.017 ± 3.01e-03 | 0.013 ± 2.15e-03 | 1.19 | 0.330 |
| C10H21O+ | 157.16 | Decanal | 0.026 ± 7.25e-03 | 0.025 ± 0.011 | 0.027 ± 0.014 | 0.021 ± 9.73e-03 | 0.06 | 0.983 |
| C10H13O2+ | 165.09 | Phenethyl acetate | 0.013 ± 2.71e-03 | 0.01 ± 9.98e-04 | 0.014 ± 4.25e-03 | 0.015 ± 3.96e-03 | 0.74 | 0.538 |
| C11H17O+ | 165.13 | Jasmone | 3.29e-03 ± 4.50e-04 | 4.98e-03 ± 8.14e-04 | 5.85e-03 ± 1.01e-03 | 4.72e-03 ± 7.42e-04 | 2.36 | 0.092 |
| C12H21O2+ | 197.15 | Geranyl acetate | 5.71e-03 ± 7.57e-04 | 5.29e-03 ± 9.65e-04 | 5.86e-03 ± 1.60e-03 | 6.81e-03 ± 1.53e-03 | 0.2 | 0.892 |
| C15H25+ | 205.20 | SQT | 0.071 ± 7.41e-03 ab | 0.046 ± 5.77e-03 a | 0.989 ± 0.184 c | 0.632 ± 0.207 bc | 11.73 | < 0.001 |
| C16H27+ | 219.21 | TMTT | 4.57e-03 ± 6.28e-04 a | 4.21e-03 ± 8.11e-04 a | 0.015 ± 2.28e-03 b | 9.51e-03 ± 1.52e-03 b | 8.22 | < 0.001 |
| C15H27O+ | 223.21 | SQT alcohols | 4.57e-03 ± 2.79e-03 ab | 1.45e-03 ± 1.05e-04 a | 9.36e-03 ± 1.52e-03 b | 0.011 ± 4.98e-03 ab | 9.21 | < 0.001 |

**Table S6.** Concentrations of volatile compounds (protonated ions) detected by PTR-TOF (Vocus S) from outdoor open-air measurements of maize plants (var. *Delprim*). Concentrations are shown as mean ± s.e. (ppb) for undamaged plants (n = 77) and plants damaged for 24 h by *Spodoptera exigua* larvae (n = 81). Measurements were conducted at two sampling distances from the plant (1–2 cm and 4–5 cm), with values correspond to concentrations averaged over 60-s measurements (1 spectrum s⁻¹). Reported ions represent protonated volatile compounds (parent ions) or fragment ions. The percentage change in volatile emissions in response to herbivory is indicated, together with W statistics and p-values from Wilcoxon rank-sum tests.

| Ions | m/z | Attribution | 1–2 cm |  |  |  |  | 4–5 cm |  |  |  |  |
| --- | --- | --- | --- | --- | --- | --- | --- | --- | --- | --- | --- | --- |
|  |  |  | Undamaged | Damaged | Change | W | P-value | Undamaged | Damaged | Change | W | P-value |
| C3H5O+ | 57.03 | Alcohols + ald. frag. | 0.421 ± 6.00e-03 | 0.421 ± 6.12e-03 | 0% | 3142 | 0.936 | 0.413 ± 5.88e-03 | 0.413 ± 5.87e-03 | 0% | 3155 | 0.9 |
| C5H7+ | 67.05 | MT frag. | 0.078 ± 1.50e-03 | 0.082 ± 1.55e-03 | 5% | 2542 | **0.045** | 0.075 ± 1.52e-03 | 0.076 ± 1.35e-03 | 1% | 3029 | 0.757 |
| C4H8N+ | 70.06 | Isobutyronitrile | 0.015 ± 3.80e-04 | 0.016 ± 4.10e-04 | 7% | 2757 | 0.209 | 0.015 ± 4.01e-04 | 0.014 ± 3.78e-04 | -7% | 3103 | 0.958 |
| C6H9+ | 81.07 | Terpenes + hexenal frag. | 0.162 ± 5.00e-03 | 0.198 ± 0.01 | 22% | 2177 | **< 0.01** | 0.148 ± 5.27e-03 | 0.153 ± 4.31e-03 | 3% | 2757 | 0.209 |
| C6H11+ | 83.09 | GLVs frag. | 0.094 ± 4.90e-03 | 0.097 ± 4.66e-03 | 3% | 2970 | 0.607 | 0.086 ± 4.70e-03 | 0.087 ± 4.47e-03 | 1% | 2912 | 0.474 |
| C5H10N+ | 84.08 | Methylbutyronitrile | 7.54e-03 ± 2.90e-04 | 7.63e-03 ± 2.69e-04 | 1% | 3015 | 0.72 | 7.16e-03 ± 2.76e-04 | 7.14e-03 ± 2.67e-04 | 0% | 3155 | 0.9 |
| C6H13+ | 85.10 | GLVs frag. | 5.75e-03 ± 1.84e-04 | 5.80e-03 ± 1.70e-04 | 1% | 3028 | 0.754 | 5.42e-03 ± 1.85e-04 | 5.48e-03 ± 1.68e-04 | 1% | 2974 | 0.616 |
| C4H10NO+ | 88.08 | Isobutyraldoxime | 6.37e-03 ± 1.74e-04 | 0.011 ± 9.17e-04 | 73% | 1198 | **< 0.001** | 6.06e-03 ± 1.27e-04 | 6.79e-03 ± 1.45e-04 | 12% | 2016 | **< 0.001** |
| C7H9+ | 93.07 | MT frag. | 0.133 ± 6.79e-03 | 0.133 ± 6.24e-03 | 0% | 3065 | 0.854 | 0.127 ± 5.93e-03 | 0.127 ± 6.25e-03 | 0% | 3212 | 0.746 |
| C7H11+ | 95.09 | Terpenes frag. | 0.063 ± 1.62e-03 | 0.08 ± 3.08e-03 | 27% | 1742 | **< 0.001** | 0.061 ± 1.65e-03 | 0.065 ± 1.51e-03 | 7% | 2432 | **0.017** |
| C6H11O+ | 99.08 | Hexenal | 0.043 ± 5.09e-04 | 0.044 ± 6.29e-04 | 2% | 2799 | 0.267 | 0.041 ± 4.41e-04 | 0.042 ± 4.92e-04 | 2% | 2835 | 0.325 |
| C6H13O+ | 101.10 | Hexenol, hexanal | 0.016 ± 4.88e-04 | 0.016 ± 4.66e-04 | 0% | 2996 | 0.671 | 0.015 ± 4.50e-04 | 0.015 ± 4.44e-04 | 0% | 3053 | 0.821 |
| C5H12NO+ | 102.09 | Methylbutanal oxime | 0.016 ± 1.31e-03 | 0.015 ± 1.32e-03 | -6% | 3201 | 0.775 | 0.012 ± 8.94e-04 | 0.011 ± 8.33e-04 | -8% | 3460 | 0.236 |
| C6H15O+ | 103.11 | Hexanol | 7.82e-04 ± 4.52e-05 | 7.64e-04 ± 4.51e-05 | -2% | 3391 | 0.344 | 6.44e-04 ± 2.70e-05 | 6.07e-04 ± 2.56e-05 | -6% | 3435 | 0.272 |
| C4H10NO2+ | 104.07 | 2-Methyl-2-nitropropane | 3.51e-03 ± 5.06e-05 | 3.67e-03 ± 5.83e-05 | 5% | 2506 | **0.033** | 3.48e-03 ± 4.89e-05 | 3.52e-03 ± 5.09e-05 | 1% | 2877 | 0.402 |
| C8H13+ | 109.10 | SQT frag. | 0.062 ± 3.29e-03 | 0.075 ± 3.56e-03 | 21% | 2300 | **< 0.01** | 0.057 ± 3.20e-03 | 0.061 ± 2.98e-03 | 7% | 2663 | 0.113 |
| C8H8N+ | 118.07 | Indole | 7.22e-03 ± 3.92e-04 | 0.048 ± 5.11e-03 | 565% | 222 | **< 0.001** | 5.54e-03 ± 2.32e-04 | 0.028 ± 2.19e-03 | 405% | 163 | **< 0.001** |
| C10H15+ | 135.12 | SQT frag. | 0.012 ± 5.13e-04 | 0.018 ± 9.43e-04 | 50% | 1653 | **< 0.001** | 0.012 ± 4.77e-04 | 0.014 ± 4.81e-04 | 17% | 2226 | **< 0.01** |
| C10H17+ | 137.13 | MT | 0.043 ± 2.14e-03 | 0.057 ± 4.62e-03 | 33% | 2250 | **< 0.01** | 0.041 ± 2.55e-03 | 0.041 ± 1.84e-03 | 0% | 2949 | 0.557 |
| C8H15O2+ | 143.11 | Hexenyl acetate | 0.02 ± 6.96e-04 | 0.022 ± 7.18e-04 | 10% | 2622 | 0.084 | 0.019 ± 7.03e-04 | 0.02 ± 6.22e-04 | 5% | 2692 | 0.138 |
| C8H17O2+ | 145.12 | Hexyl acetate | 3.65e-03 ± 1.59e-04 | 3.64e-03 ± 1.40e-04 | 0% | 3076 | 0.884 | 3.56e-03 ± 1.58e-04 | 3.57e-03 ± 1.40e-04 | 0% | 2985 | 0.644 |
| C11H17+ | 149.13 | SQT frag. | 4.04e-03 ± 1.57e-04 | 0.015 ± 1.31e-03 | 271% | 307 | **< 0.001** | 3.83e-03 ± 1.48e-04 | 7.09e-03 ± 2.80e-04 | 85% | 681 | **< 0.001** |
| C9H11O2+ | 151.08 | Phenylmethyl acetate | 6.01e-03 ± 1.33e-04 | 6.07e-03 ± 1.30e-04 | 1% | 2972 | 0.612 | 5.97e-03 ± 1.30e-04 | 5.92e-03 ± 1.26e-04 | -1% | 3199 | 0.781 |
| C11H19+ | 151.15 | DMNT | 2.82e-03 ± 1.01e-04 | 6.65e-03 ± 4.72e-04 | 136% | 490 | **< 0.001** | 2.72e-03 ± 1.00e-04 | 3.96e-03 ± 1.61e-04 | 46% | 1300 | **< 0.001** |
| C10H21O+ | 157.16 | Decanal | 0.013 ± 9.13e-04 | 0.013 ± 8.75e-04 | 0% | 3067 | 0.859 | 0.012 ± 8.64e-04 | 0.012 ± 8.25e-04 | 0% | 2980 | 0.631 |
| C10H13O2+ | 165.09 | Phenethyl acetate | 3.87e-03 ± 8.66e-05 | 3.78e-03 ± 8.13e-05 | -2% | 3413 | 0.306 | 3.83e-03 ± 8.31e-05 | 3.77e-03 ± 7.93e-05 | -2% | 3349 | 0.424 |
| C11H17O+ | 165.13 | (Z)-jasmone | 1.34e-03 ± 4.24e-05 | 1.55e-03 ± 4.44e-05 | 16% | 2108 | **< 0.001** | 1.28e-03 ± 4.08e-05 | 1.39e-03 ± 3.38e-05 | 9% | 2311 | **< 0.01** |
| C12H21O2+ | 197.15 | Geranyl acetate | 1.45e-03 ± 4.96e-05 | 1.46e-03 ± 4.75e-05 | 1% | 3033 | 0.767 | 1.47e-03 ± 5.15e-05 | 1.44e-03 ± 4.85e-05 | -2% | 3267 | 0.607 |
| C15H25+ | 205.20 | SQT | 3.34e-03 ± 1.86e-04 | 0.038 ± 4.38e-03 | 1038% | 66 | **< 0.001** | 2.98e-03 ± 1.76e-04 | 0.013 ± 8.90e-04 | 336% | 158 | **< 0.001** |
| C16H27+ | 219.21 | TMTT | 1.26e-03 ± 8.19e-05 | 1.41e-03 ± 7.36e-05 | 12% | 2491 | **0.029** | 1.23e-03 ± 8.24e-05 | 1.32e-03 ± 7.14e-05 | 7% | 2658 | 0.11 |
| C15H27O+ | 223.21 | SQT alcohols | 4.96e-04 ± 2.17e-05 | 6.80e-04 ± 3.16e-05 | 37% | 1857 | **< 0.001** | 5.06e-04 ± 2.61e-05 | 5.56e-04 ± 2.37e-05 | 10% | 2586 | 0.064 |

**Table S7.** Median concentrations of selected herbivory-associated volatiles measured during laboratory assays using dynamic headspace sampling and during open-air measurements at two sampling distances, with fold-change comparisons.

| Compound | Lab (ppb) | Open air<br>1–2 cm (ppb) | Open air<br>4–5 cm (ppb) | Fold change<br>(Lab → 1–2 cm) | Fold change<br>(Lab → 4–5 cm) | Fold change<br>(1–2 → 4–5 cm) |
| --- | --- | --- | --- | --- | --- | --- |
| C<sub>8</sub>H<sub>8</sub>N<sup>+</sup> | 6.6 | 0.030 | 0.023 | –220× | –287× | –1.3× |
| C<sub>15</sub>H<sub>25</sub><sup>+</sup> | 0.60 | 0.023 | 0.012 | –26× | –50× | –1.9× |

**Table S8.** Performance of machine-learning classifiers evaluated using K-fold cross-validation. Mean classification accuracy ± standard deviation across five folds is reported for the training datasets. Individual maize plants were sampled outdoors for 60 s at distances of 1–2 cm and 4–5 cm using PTR-TOF (Vocus S). The 60-s measurements were segmented into intervals of 30, 15, 5, and 1 s, with signals averaged within each interval, to assess the effect of sampling duration on prediction performance. Two validation strategies were applied. In the first approach, all time intervals were treated as independent observations using a grouped K-fold cross-validation scheme to account for plant identity. In the second approach, to maintain one observation per plant, only the first interval of each duration was retained for model training and evaluation.

| Dataset | Classifier | 1–2 cm, all intervals |  | 4–5 cm, all intervals |  | 1–2 cm, first interval |  | 4–5 cm, first interval |  |
| --- | --- | --- | --- | --- | --- | --- | --- | --- | --- |
|  |  | Acc. mean | 95% CI | Acc. mean | 95% CI | Acc. mean | 95% CI | Acc. mean | 95% CI |
| 1 second | Logistic Regression | 0.9 | 0.04 | 0.89 | 0.05 | 0.85 | 0.05 | 0.83 | 0.08 |
|  | KNeighbors | 0.69 | 0.1 | 0.71 | 0.04 | 0.6 | 0.09 | 0.58 | 0.11 |
|  | SVM | 0.84 | 0.06 | 0.86 | 0.04 | 0.73 | 0.08 | 0.65 | 0.08 |
|  | Decision Tree | 0.84 | 0.05 | 0.85 | 0.05 | 0.75 | 0.21 | 0.76 | 0.06 |
|  | Random Forest | 0.89 | 0.05 | 0.89 | 0.03 | 0.85 | 0.06 | 0.87 | 0.06 |
| 5 seconds | Logistic Regression | 0.91 | 0.05 | 0.9 | 0.06 | 0.8 | 0.05 | 0.89 | 0.05 |
|  | KNeighbors | 0.7 | 0.13 | 0.71 | 0.02 | 0.65 | 0.05 | 0.65 | 0.15 |
|  | SVM | 0.83 | 0.08 | 0.84 | 0.06 | 0.75 | 0.09 | 0.73 | 0.15 |
|  | Decision Tree | 0.87 | 0.06 | 0.88 | 0.07 | 0.81 | 0.09 | 0.84 | 0.05 |
|  | Random Forest | 0.92 | 0.04 | 0.9 | 0.05 | 0.9 | 0.1 | 0.9 | 0.05 |
| 15 seconds | Logistic Regression | 0.91 | 0.05 | 0.92 | 0.06 | 0.89 | 0.08 | 0.9 | 0.07 |
|  | KNeighbors | 0.7 | 0.13 | 0.73 | 0.09 | 0.65 | 0.19 | 0.68 | 0.15 |
|  | SVM | 0.81 | 0.07 | 0.83 | 0.08 | 0.73 | 0.09 | 0.74 | 0.16 |
|  | Decision Tree | 0.87 | 0.06 | 0.9 | 0.07 | 0.86 | 0.08 | 0.8 | 0.1 |
|  | Random Forest | 0.94 | 0.05 | 0.92 | 0.04 | 0.92 | 0.07 | 0.91 | 0.06 |
| 30 seconds | Logistic Regression | 0.92 | 0.06 | 0.94 | 0.05 | 0.94 | 0.06 | 0.94 | 0.09 |
|  | KNeighbors | 0.7 | 0.14 | 0.72 | 0.07 | 0.6 | 0.17 | 0.65 | 0.09 |
|  | SVM | 0.81 | 0.07 | 0.82 | 0.08 | 0.71 | 0.2 | 0.8 | 0.19 |
|  | Decision Tree | 0.9 | 0.1 | 0.93 | 0.03 | 0.86 | 0.07 | 0.85 | 0.05 |
|  | Random Forest | 0.94 | 0.07 | 0.94 | 0.04 | 0.93 | 0.05 | 0.93 | 0.05 |
| 60 seconds | Logistic Regression | 0.94 | 0.05 | 0.96 | 0.05 | 0.93 | 0.09 | 0.96 | 0.05 |
|  | KNeighbors | 0.7 | 0.15 | 0.75 | 0.03 | 0.66 | 0.09 | 0.75 | 0.09 |
|  | SVM | 0.84 | 0.06 | 0.83 | 0.09 | 0.77 | 0.14 | 0.85 | 0.09 |
|  | Decision Tree | 0.93 | 0.03 | 0.85 | 0.11 | 0.9 | 0.03 | 0.89 | 0.09 |
|  | Random Forest | 0.95 | 0.06 | 0.9 | 0.06 | 0.95 | 0.04 | 0.9 | 0.1 |

**Table S9.**
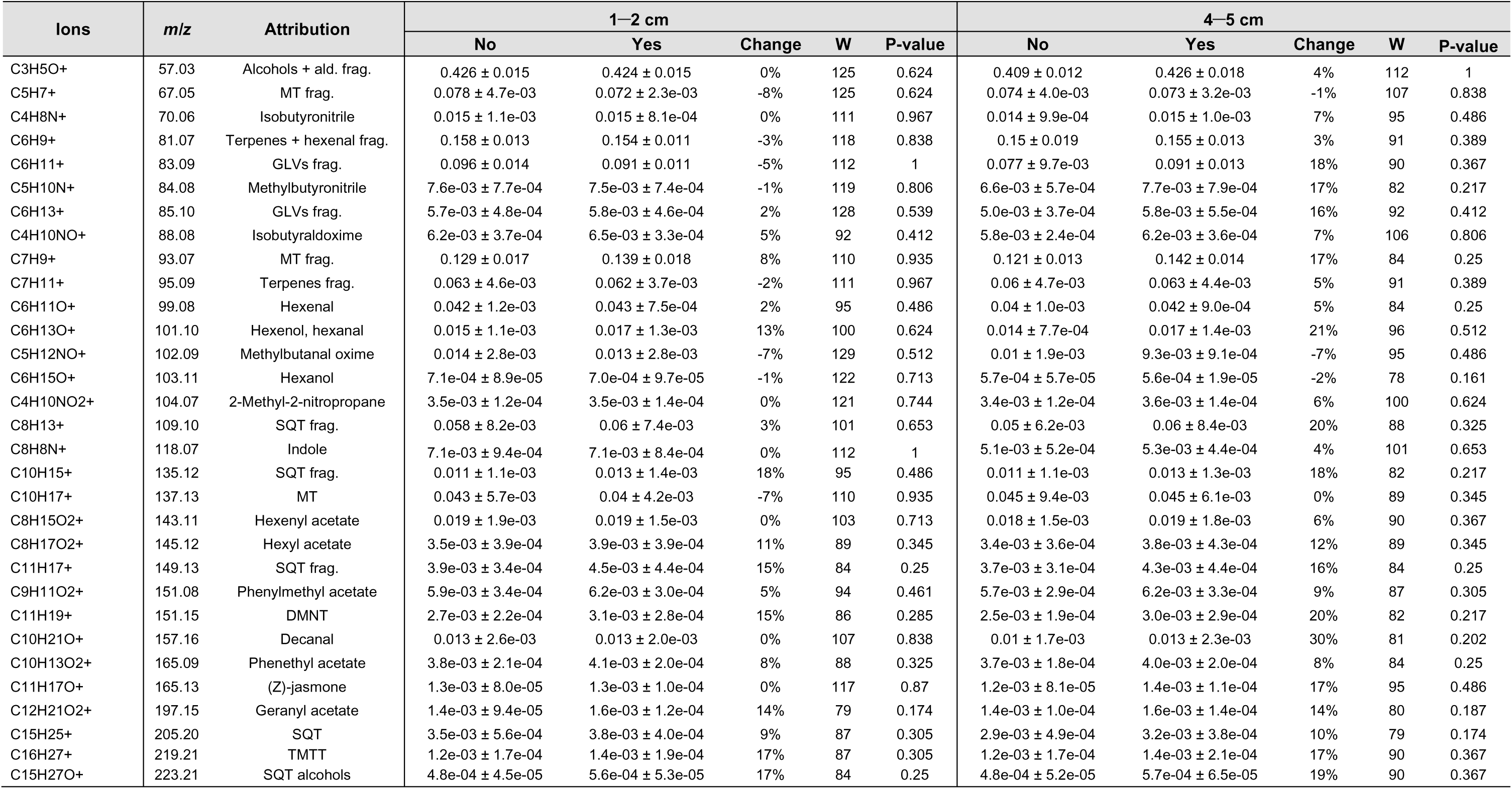
Effect of bagging maize plants in PET cooking bags on volatile emissions detected by PTR-TOF (Vocus S) from outdoor open-air measurements. Concentrations are shown as mean ± s.e. (ppb) for unbagged (“No”) and bagged (“Yes”) plants. Measurements were conducted at two sampling distances from the plant (1–2 cm and 4–5 cm), with values correspond to concentrations averaged over 60-s measurements (1 spectrum s⁻¹). Reported ions represent protonated volatile compounds (parent ions) or fragment ions. To assess whether bagging itself induced volatile emissions, 25% of control plants (five per sampling day) were bagged using the same procedure applied to damaged plants and compared with five unbagged control plants randomly selected on the same day to ensure comparable sample sizes. The percentage change in volatile emissions due to bagging is reported, together with W statistics and p-values from Wilcoxon rank-sum tests.

